# Non-invasive temporal interference stimulation targeting the hippocampus enhances fast motor memory consolidation

**DOI:** 10.64898/2026.11.11.766003

**Authors:** Xiaoxi Pan, Jet Taylor, Ines R. Violante, Nir Grossman, Cory Inman, Bradley R. King, Geneviève Albouy

**Affiliations:** Department of Health and Kinesiology, College of Health, University of Utah, Salt Lake City, USA; School of Biomedical Engineering and Imaging Sciences, King’s College London, London, UK; Division of Brain Sciences, Imperial College London, London, UK; Department of Psychology, University of Utah, Salt Lake City, USA

**Keywords:** Motor Sequence Learning, Memory Consolidation, Hippocampus, Temporal Interference Stimulation

## Abstract

Converging neuroimaging evidence indicates that the hippocampus is involved in motor sequence learning (MSL) and its consolidation at multiple timescales. However, direct evidence for a causal role of the hippocampus in these processes in healthy humans remains scarce, partly due to the inability of conventional non-invasive brain stimulation techniques to reach this deep brain structure. Here, we used temporal interference stimulation (TIS) — a technique enabling non-invasive stimulation targeted toward deep brain regions — to probe the functional role of the hippocampus in MSL and its consolidation. In two separate experiments, sixty-four young healthy adults (36 females) received, in within-subject counterbalanced designs, active (5 Hz envelope) or control (0 Hz envelope) hippocampal-targeted TIS either during active practice of an MSL task (task-TIS experiment, n = 32) or during the short rest periods interleaved with task practice known to support rapid forms of consolidation (rest-TIS experiment, n = 32). Consolidation was assessed at both the micro timescale (i.e., over the short “micro-offline” rest intervals interspersed with task practice) and macro timescale (i.e., over a 24-hour “macro-offline” retention interval). Our results indicate that TIS targeting the hippocampus during inter-practice rest periods — but not during task practice — significantly enhanced micro-offline gains in performance as compared to control stimulation. While this effect did not carry over to the 24-hour retention interval at the group level, participants with greater stimulation-induced micro-offline gains showed larger overnight performance improvements. These findings provide the first direct evidence that experimental interventions targeting the hippocampus modulate offline motor memory consolidation processes.

## Introduction

A large body of neuroimaging evidence indicates that the hippocampus is recruited during motor sequence learning (MSL) irrespective of whether learning is explicit or implicit in nature (Albouy et al., 2008, 2012; Albouy, King, et al., 2013; Fernández-Seara et al., 2009; Gheysen et al., 2010; King et al., 2022; Pinsard et al., 2019; Schendan et al., 2003). Previous studies using both univariate and multivariate analyses of functional magnetic resonance imaging (fMRI) data suggest that the hippocampus supports the formation of spatio-temporal representations of motor sequences that capture the goal of the sequence of movements to be performed (Albouy et al., 2015; Albouy, Fogel, et al., 2013; Dolfen et al., 2024). Interestingly, these hippocampal representations have been associated with motor memory consolidation, i.e., the offline process by which memory traces are reorganized into more robust representations in the absence of further task practice (Dudai et al., 2015; Frankland & Bontempi, 2005), and this involvement was observed at different timescales. At the micro timescale (seconds to minutes), the short rest intervals interspersed with task practice known to support rapid forms of motor memory consolidation (Bönstrup et al., 2019, 2020) are accompanied by increased hippocampal activity (Gann et al., 2023; Jacobacci et al., 2020), elevated hippocampal ripple rate (Chen et al., 2025; Sjøgård et al., 2025) and hippocampo-neocortical replay of the learned motor sequence (Buch et al., 2021). At the macro timescale (hours to days), multivoxel patterns elicited in the hippocampus during MSL have been shown to persist into subsequent periods of awake rest and reflect the spontaneous reactivation of the spatiotemporal representations of the learned motor sequences (King et al., 2022). For timescales spanning a night of sleep, earlier neuroimaging studies demonstrated that hippocampal activity and connectivity patterns during initial MSL predict performance improvement after a night of sleep (Albouy et al., 2008; Albouy, Sterpenich, et al., 2013). It was also suggested that the beneficial effect of sleep on motor memory consolidation was preferentially observed for the spatial, hippocampal-mediated representations of MSL (Albouy et al., 2015; Albouy, Fogel, et al., 2013).

Altogether, the evidence reviewed above challenges traditional models of memory organization (Cohen & Squire, 1980; Squire & Zola, 1996) as it suggests that the hippocampus plays a critical role in motor sequence memory consolidation at both the micro and macro timescales. However, these previous neuroimaging studies are correlational in nature, and causal evidence remains limited, in part because conventional non-invasive brain stimulation techniques cannot directly reach this deep brain structure. In this study pre-registered in the Open Science Framework (https://osf.io/5m37g), we leveraged recent advances in transcranial electrical stimulation and used temporal interference stimulation (TIS), a novel non-invasive brain stimulation technique (Grossman et al., 2017) that allows targeted stimulation of deep brain structures in humans <u>(e.g., Beanato et al., 2024; Violante et al., 2023; Wessel et al., 2023)</u>, to test whether stimulation targeting the hippocampus modulates MSL and its consolidation. To do so, active (5 Hz envelope) and control (0 Hz envelope) TIS were applied in sixty-four young healthy adults (36 females) during MSL, in two experiments using within-subject counterbalanced designs (Figure 1A), with a montage previously validated to modulate hippocampal function (Figure 1C; Beanato et al., 2024; Violante et al., 2023). In these two separate experiments (n = 32 in each experiment), stimulation was delivered either during task practice (referred to as the task-TIS experiment) or during the short rest periods interspersed with task practice that support the rapid consolidation process (rest-TIS experiment, see Figure 1A). Consolidation was assessed at the micro timescale as the changes in performance over the inter-practice rest periods and at the macro timescale as the changes in performance over a 24-hour interval including a night of sleep (Figure 1B). Based on previous work suggesting that the hippocampus encodes spatiotemporal representations of the learned motor sequences (Albouy et al., 2015; Albouy, Fogel, et al., 2013; Dolfen et al., 2024), we also implemented a representation test similar to our previous research (Albouy et al., 2015; Albouy, Fogel, et al., 2013) to examine the effect of hippocampal-targeted TIS on spatial (as compared to motoric) representations of the task (see Figure 1A). Given the previous evidence reviewed above suggesting that the hippocampus is specifically linked to offline motor memory consolidation processes (Albouy et al., 2008; Albouy, King, et al., 2013; Buch et al., 2021; Jacobacci et al., 2020; Mylonas et al., 2024), we hypothesized that active TIS, as compared to control TIS, targeting the hippocampus would preferentially enhance offline motor memory consolidation at both the micro and macro timescales, especially when applied during rest periods (Buch et al., 2021; Gann et al., 2023; Jacobacci et al., 2020; Mylonas et al., 2024). We further predicted that hippocampal-targeted TIS would enhance the spatial representation of the MSL task.

**Figure 1.**
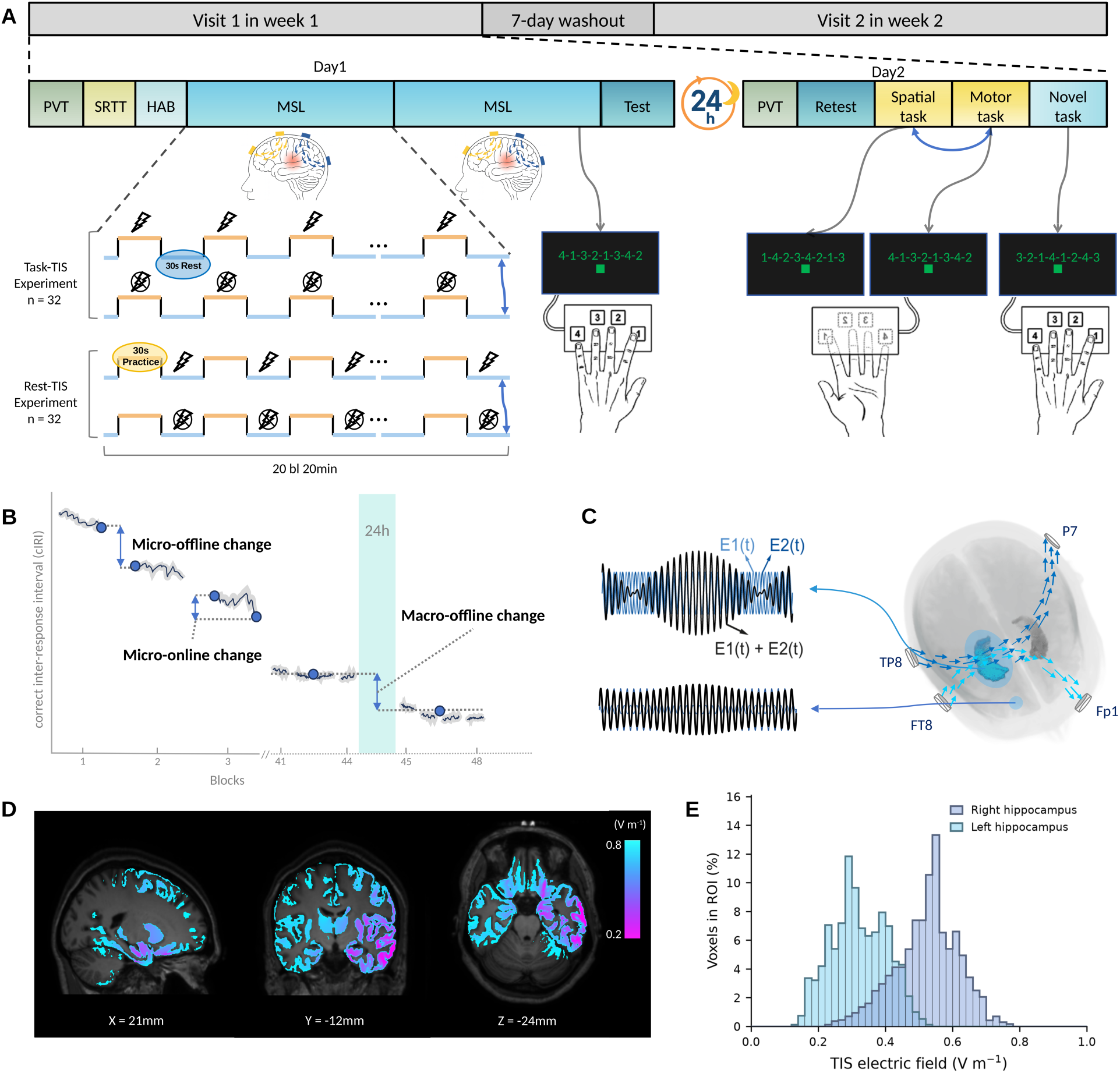
Experimental procedure and Temporal Interference Stimulation (TIS) principle. A. Experimental procedure. Sixty-four young healthy adults were randomly distributed in one of two experiments (task-TIS and rest-TIS) that included two visits (Visit 1 on week 1 and Visit 2 on week 2), each consisting of two consecutive testing days (Day 1 and Day 2). Participants followed a regular sleep schedule from three days before Day 1 through Day 2, monitored with sleep diaries and wrist actigraphy (see Methods); sleep and vigilance did not differ between stimulation conditions and experiments (see section 1 of the Supplemental information). On Day 1, participants completed a Psychomotor Vigilance Task (PVT) to assess vigilance, a random Serial Reaction Time Task (SRTT) to assess baseline motor execution, and a TIS habituation stimulation phase (HAB, see Methods). They then performed two 20-block (bl) training sessions (Training 1 and Training 2) on the Motor Sequence Learning (MSL) task during which task practice (30-s practice blocks) alternated with inter-practice rest periods (30-s rest blocks). During training, either active (lightning bolt) or control (crossed lightning bolt) stimulation targeting the hippocampus was applied during the different visits (order of the stimulation conditions counterbalanced across participants). In the task-TIS experiment (n = 32), hippocampal-targeted TIS was delivered during the 30-s practice periods, while in the rest-TIS experiment (n = 32) it was applied during the 30-s inter-practice rest periods. The MSL task involved repeatedly tapping a visually presented 8-digit finger sequence (e.g. 4-1-3-2-1-3-4-2, with 1 = index finger and 4 = little finger) using the non-dominant left hand on a custom keyboard. A stimulation- and fatigue-free post-training test (4 blocks) followed the training phases. On Day 2, participants were retested on the learned sequence (4 blocks) and on the spatial and motor representations of the learned sequence (order of representation tests counterbalanced across participants) using an inverted keyboard (see Methods). The spatial task preserved the spatial layout of the trained sequence but required different finger movements (e.g. 1-4-2-3-4-2-1-3), while the motor task preserved the finger movement sequence but altered the spatial layout (e.g. 4-1-3-2-1-3-4-2). Finally, a novel sequence task was performed to assess learning-specificity (e.g. 3-2-1-4-1-2-4-3). Note that different motor sequences were learned during the two different visits (see Methods). **B. Assessment of consolidation at the micro and macro timescales.** Schematic illustration of the performance speed measure (mean correct inter-response interval, cIRI) across practice blocks on Day 1 and at the Day 2 retest. Micro-online changes in performance were computed as the difference in mean cIRI between the first and the last eight correct transitions within a practice block and micro-offline changes as the difference between the last eight correct transitions of a block and the first eight of the subsequent block (mean cIRI depicted as the blue dots in blocks 1-3). Macro-offline changes in performance were computed as the difference in mean cIRI between the average performance across the four post-training test blocks on Day 1 and the four retest blocks on Day 2 (i.e., across the 24-hour interval, mean cIRI depicted as the blue dots in blocks 41-48). **C. TIS principle and electrode montage.** TIS targeted the right hippocampus using a montage validated in previous research (Beanato et al., 2024; Violante et al., 2023). Two pairs of scalp electrodes placed at FT8-Fp1 and TP8-P7 according to the EEG 10-10 system delivered two sinusoidal high-frequency currents at 2000 Hz and 2005 Hz at a 2 mA intensity (baseline-to-peak) per channel. The resulting fields E₁(t) and E₂(t) superimpose within the brain, yielding an envelope modulated at the difference frequency (Δf = 5 Hz) that can be targeted toward deep structures, here the right hippocampus (see panels D-E for electric field simulations). High-frequency control stimulation was applied using the same montage with both channels delivering a 2000 Hz current, producing a flat envelope. Stimulation-related sensations were collected with standardized questionnaires during HAB and after each training session, and a blinding questionnaire was administered at the end of each Day 1 visit (see panel A). The results of the corresponding exploratory analyses are reported in section 2 of the Supplemental information. Briefly, results indicate that (i) TIS was well tolerated, (ii) no unanticipated or serious adverse events occurred, (iii) sensations did not differ between active and control stimulation conditions, and that (iv) participants remained blind to the stimulation condition. **D. Electric field modeling.** The TIS-induced electric fields were simulated in exploratory analyses performed using the MIDA anatomical head model (Iacono et al., 2015) with the experimental montage and current intensities mentioned above with anisotropic conductivity in white matter and midbrain derived from diffusion MRI (see Methods). Colors depict the TIS electric field in grey matter tissues, quantified as the amplitude of the low-frequency envelope modulation of the field (V m⁻¹), on sagittal, coronal, and axial sections through the right hippocampus (MNI coordinates below each section). Values below 0.2 V m⁻¹ are not displayed. **E. TIS electric field distribution in the right and left hippocampi.** Histograms show the field distribution across voxels of each hippocampus, expressed as the percentage of voxels (right: dark blue; left: light blue; see Methods), indicating preferential exposure of the targeted right hippocampus (median field strength in V m⁻¹; right: 0.53, left: 0.31). See also section 3 in the Supplemental information for a depiction of field distribution in other regions of interest including the putamen (median field strength in V m⁻¹; right: 0.49, left 0.33), caudate (right: 0.38, left: 0.35), thalamus (right: 0.37, left: 0.30) and M1 (right: 0.36, left: 0.22); and see discussion for focality considerations and how potential field exposure in the putamen might have influenced the current pattern of results.

## Results

### Hippocampal-targeted TIS did not affect overall motor performance during initial learning

We first tested whether hippocampal-targeted TIS (compared to control TIS) modulated overall motor performance speed during initial MSL on Day 1 of each visit. Results of both experiments indicate that active TIS, irrespective of whether it was applied during task practice (Figure 2A) or during inter-practice rest intervals (Figure 2B), did not influence motor performance as compared to control stimulation (repeated-measures ANOVAs using block [40 for training and 4 for test] and stimulation conditions [active vs. control] as within-subject factors; task-TIS experiment: no main effect of stimulation condition during training, F(1, 31) = 0.15, p = .697, ηp² = .005 and test, F(1, 31) = 0.01, p = .919, ηp² < .001, no block by stimulation condition during training, F(3.96, 122.84) = 1.06, p = .381, ηp² = .033 and test, F(1.88, 58.35) = 0.11, p = .883, ηp² = .004; rest-TIS experiment: no main effect stimulation condition during training, F(1, 31) = 0.04, p = .836, ηp² = .001 and test, F(1, 31) = 3.00, p = .093, ηp² = .088, no block by stimulation condition during training, F(3.99, 123.58) = 0.33, p = .856, ηp² = .011 and test, F(2.41, 74.60) = 0.15, p = .894, ηp² = .005; see section 4 in the Supplemental information for the full statistical model and for a statistical comparisons between experiments). Altogether, these results indicate that hippocampal-targeted TIS applied during task practice or inter-practice rest periods did not influence overall motor performance during initial learning (and see section 5 in the Supplemental information for similar results on performance accuracy).

**Figure 2.**
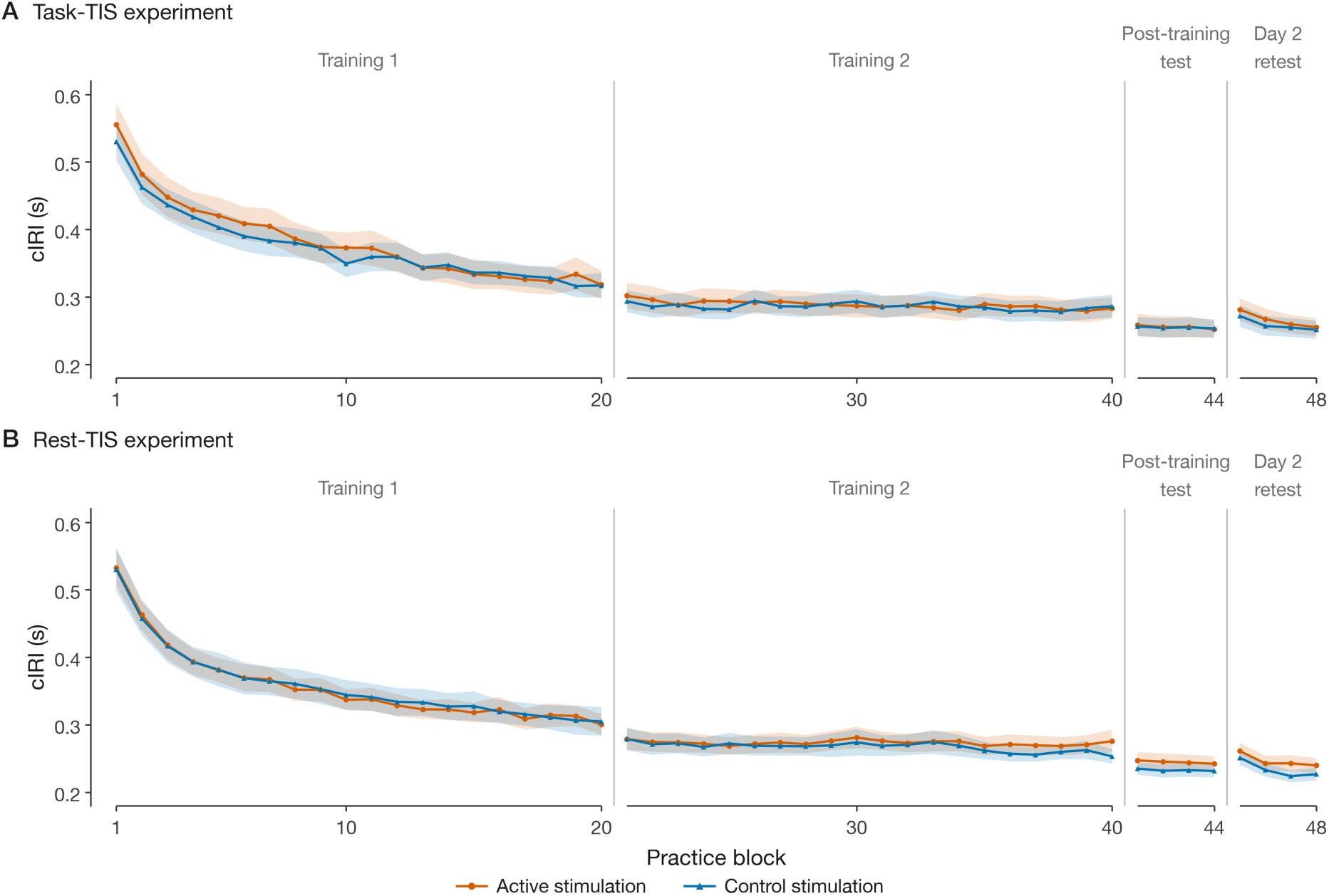
Performance speed for the task-TIS. (A) and rest-TIS (B) experiments (n = 32 in each experiment) and for the active (orange) and control (blue) stimulation conditions. Performance speed, measured as the inter-response interval for correct key presses (cIRI, in s), is plotted for each block of Training 1 (blocks 1–20), Training 2 (blocks 21– 40), Post-training test (41-44) and for the Day 2 retest (45-48). Lower values indicate faster performance. Shaded regions represent SEM. Hippocampal-targeted TIS - irrespective of whether it was applied during task practice or during inter-practice rest periods - did not influence overall motor performance. Performance on the spatial and motor representations of the task as well as on the novel sequence are presented in section 6 of the Supplemental information. Results did not show an effect of stimulation on these tasks.

### Hippocampal-targeted TIS administered during inter-practice rest periods, but not during task practice, enhanced fast motor memory consolidation

We next examined whether hippocampal-targeted TIS influenced motor memory consolidation at the micro timescale. Micro timescale changes in performance were computed across the inter-practice rest periods (micro-offline changes in performance) and over the practice periods (micro-online changes in performance, see Figure 1B and Methods for details). Figure 3A-B depicts the accumulation of both micro-offline and micro-online changes in performance across the 40 training blocks (positive/negative values indicate faster/slower performance across the interval). In both experiments, micro-offline changes were substantially larger than micro-online changes in performance (micro-learning type (2: online vs. offline) x stimulation condition (2: active vs. control); main effect of micro-learning type; task-TIS experiment: F(1, 31) = 92.50, p < .001, ηp² = .749, rest-TIS experiment: F(1, 31) = 31.68, p < .001, ηp² = .505), indicating that, as expected (Bönstrup et al., 2020; Mylonas et al., 2024), performance improvements in healthy young adults accumulated over the inter-practice rest intervals rather than over the practice period. Critically, while hippocampal-targeted TIS applied during task practice did not modulate these micro timescale metrics (task-TIS experiment; micro-learning type × stimulation condition: F(1, 31) = 0.18, p = .677, ηp² = .006, Figure 3A, 3C), stimulation administered during the inter-practice rest intervals did (rest-TIS experiment; micro-learning type × stimulation condition: F(1, 31) = 8.67, p = .006, ηp² = .219, Figure 3B, 3D). Specifically, follow-up paired-samples t-tests showed that, in the rest-TIS experiment, the sum of the micro-offline gains in performance accumulated over learning was greater in the active compared to the control stimulation condition (rest-TIS experiment; active: M = 1.51, SD = 1.33; control: M = 1.07, SD = 1.14; t(31) = 3.10, p = .004, Cohen’s dz = 0.55). Concurrently, the micro-online decay in performance was greater in the active compared to the control stimulation condition (rest-TIS experiment; active: M = −1.29, SD = 1.42; control: M = −0.82, SD = 1.12; t(31) = 2.77, p = .009, dz = 0.49). Notably, between-experiment comparisons revealed that these effects were significantly greater in the rest-TIS than in the task-TIS experiment (mixed-design ANOVA with experiment [task-TIS vs. rest-TIS] as between-subject factor and micro-learning type and stimulation condition as within-subject factors; experiment × micro-learning type × stimulation condition interaction: F(1, 62) = 5.32, p = .024, ηp² = .079). Altogether, these results indicate that hippocampal-targeted TIS applied during inter-practice rest periods, but not during task practice, enhanced micro-offline changes in performance reflecting the fast motor memory consolidation process, in parallel with a reduction in micro-online learning.

**Figure 3.**
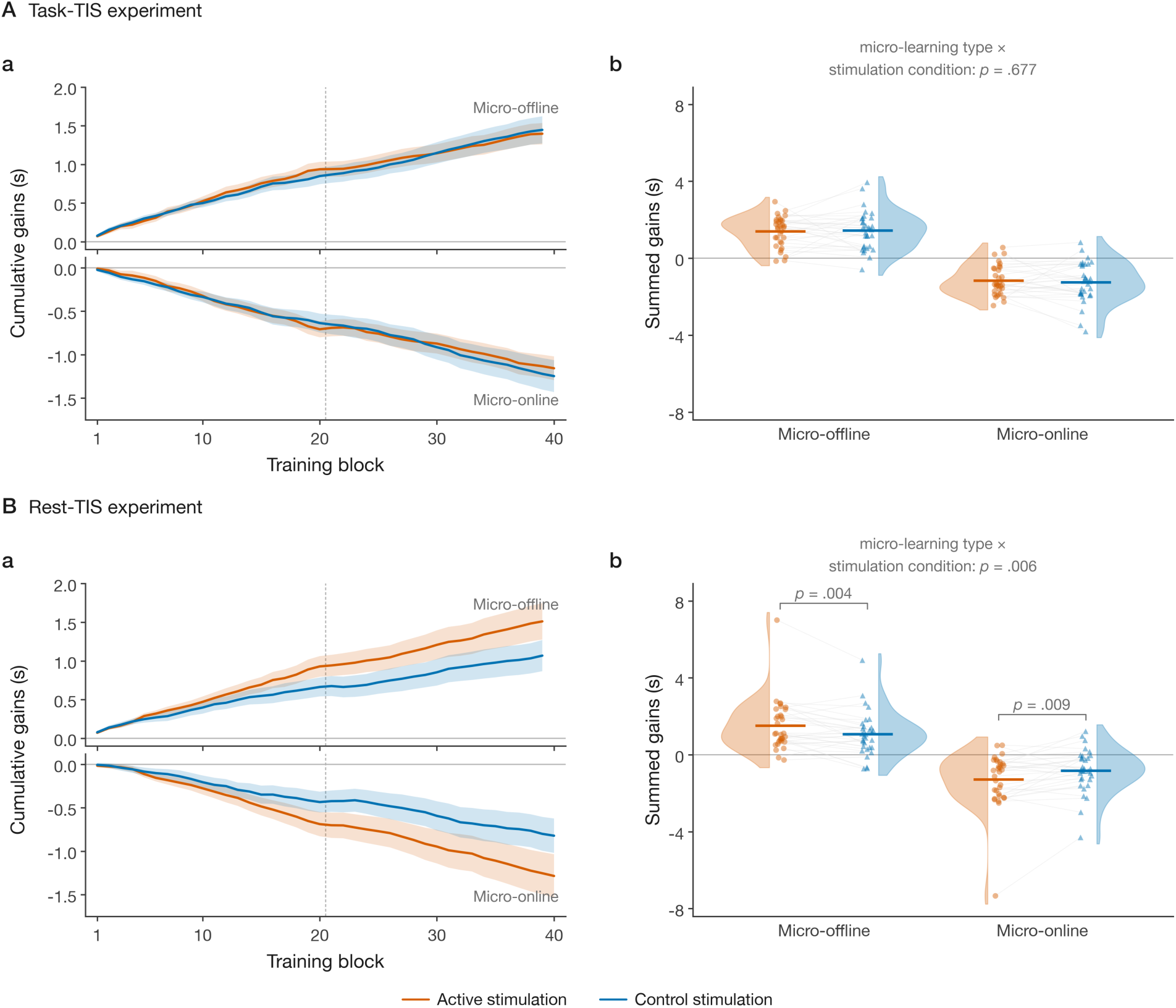
Learning and consolidation at the micro timescale for the task-TIS. (A) and rest-TIS (B) experiments (n = 32 in each experiment) and for the active (orange) and control (blue) stimulation conditions. **A. Task-TIS experiment. a.** Cumulative micro-offline (upper panel) and micro-online (lower panel) changes in performance plotted as a function of training blocks. The dashed vertical line marks the transition between Training 1 and 2. Shaded regions represent SEM. **b.** Sum of the micro-offline and micro-online changes in performance across the 40 training blocks. Colored dots depict individual data, shaded areas represent the kernel density estimate, thick horizontal lines denote group means, and grey lines connect the two stimulation conditions within participants. Hippocampal-targeted TIS applied during task practice did not influence learning and consolidation at the micro timescale. **B. Rest-TIS experiment. a.** Hippocampal-targeted TIS applied during inter-practice rest intervals enhanced micro-offline changes in performance in parallel with a reduction of micro-online learning. This stimulation effect was already significant at the end of Training 1, i.e., during the early phase of learning (blocks 1–20; see Methods and Supplemental section 7 for statistics). Note that these results remained significant when excluding the participant showing extreme micro-offline gains in performance (micro-learning type x stimulation condition interaction: F(1, 30) = 7.30, p = .011, ηp² = .196, see Supplemental section 7 for full models including or excluding this extreme participant).

### Hippocampal-targeted rest-TIS tended to enhance overnight motor memory consolidation in TIS responders

We assessed whether hippocampal-targeted TIS applied during initial MSL influenced consolidation at the macro timescale, i.e., over the 24 h consolidation interval (see Figure 1B and Figure 4A-B, in which positive values indicate improved performance). Overnight changes in performance did not differ between stimulation conditions in either experiment (paired-samples t-tests, active vs. control; task-TIS experiment: t(31) = 1.24, p = .226, dz = 0.22; rest-TIS experiment: t(31) = 0.24, p = .810, dz = 0.04; see Supplemental section 8 for comparisons between experiments). Hippocampal-targeted TIS applied during initial learning therefore did not enhance consolidation over the 24-hour retention interval. We then tested the relationship between performance changes at the micro and macro timescales (see Figure 4C-D). The correlation between micro-offline and macro-offline changes in performance was not significant in either stimulation condition in the task-TIS experiment (active: r = −.29, p = .112; control: r = −.08, p = .682; Pearson & Filon’s z testing the difference between the two conditions = −0.97, p = .334; Figure 4C). In contrast, in the rest-TIS experiment, participants who accumulated larger micro-offline gains in performance in the active stimulation condition showed greater overnight gains in performance (r = .38, p = .031, Figure 4D). This association was not significant in the control stimulation condition (r = .14, p = .442), although the correlation coefficients did not significantly differ between stimulation conditions (Pearson & Filon’s z = 1.34, p = .180). Interestingly, an exploratory correlation analysis revealed that the positive micro-macro relationship observed in the active rest-TIS experiment was significantly greater than in the active task-TIS experiment (Fisher z = −2.65, p = .008; and note that no such difference was observed for the control condition; Fisher z = −0.83, p = .408). Altogether, these results suggest that hippocampal-targeted TIS applied during inter-practice rest periods tended to enhance consolidation at the macro timescale in participants showing a greater effect of stimulation on consolidation at the micro timescale.

**Figure 4.**
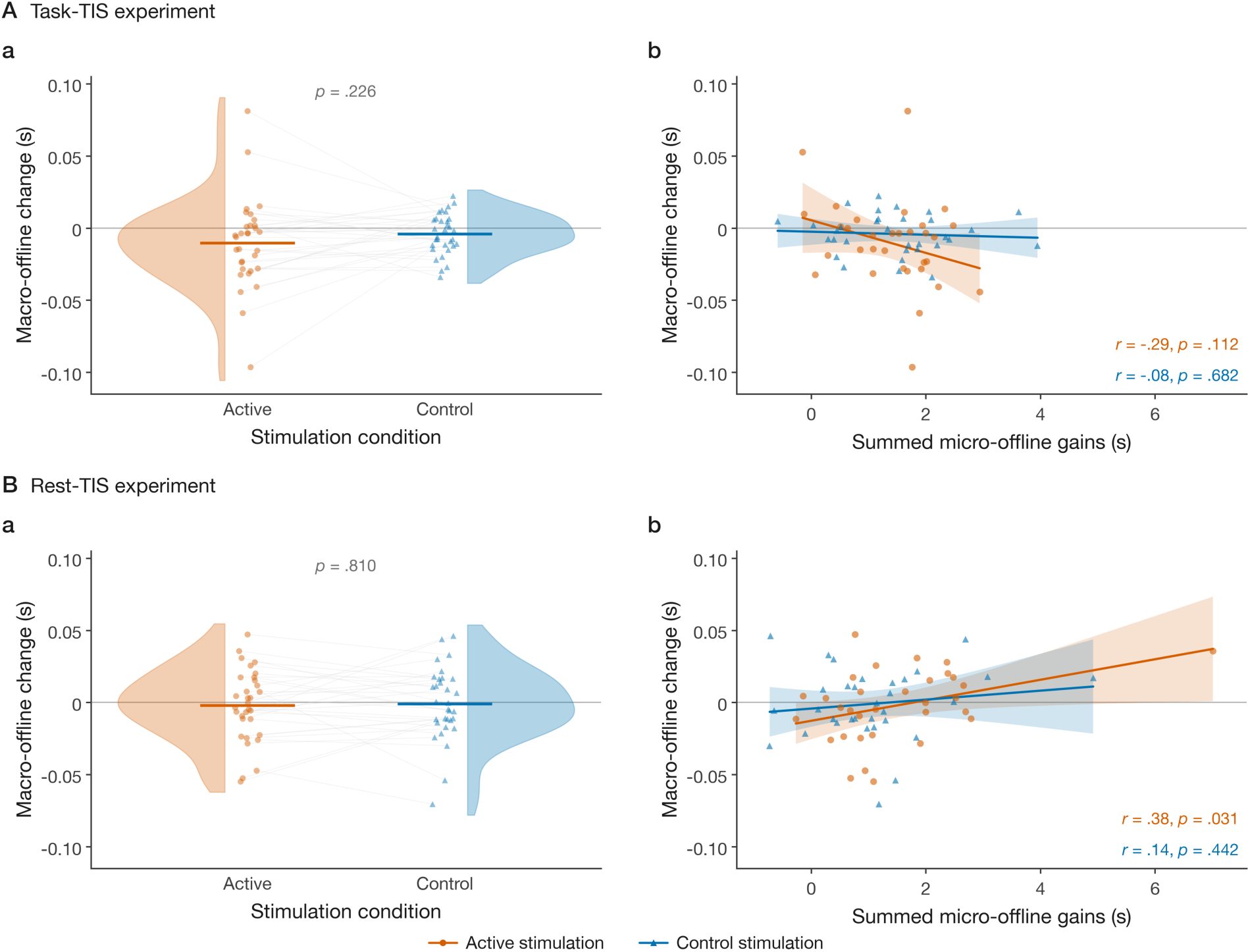
Consolidation at the macro timescale and its relationship with the micro timescale for the task-TIS. (A) and rest-TIS (B) experiments (n = 32 in each experiment) and for the active (orange) and control (blue) stimulation conditions. **A. Task-TIS experiment. a.** Macro-offline changes in performance did not differ between stimulation conditions (positive values indicate an overnight improvement in performance). Colored dots depict individual data, shaded areas represent the kernel density estimate, thick horizontal lines denote group means, and grey lines connect the two stimulation conditions within participants. **b.** The correlation between micro- and macro-offline changes in performance did not yield significant results. Dots depict individual participants, solid lines the linear fits for each stimulation condition, and shaded areas the 95% confidence intervals of the fits. **B. Rest-TIS experiment. a.** Macro-offline changes in performance did not differ between stimulation conditions. **b.** The correlation between micro- and macro-offline changes in performance showed that participants who accumulated larger micro-offline gains in the active, but not control, stimulation condition showed greater overnight performance improvements. Note that this correlation was no longer significant after excluding the participant showing extreme micro-offline gains in performance (r = .27, p = .136) although the between-experiment difference in correlations remained significant in the active stimulation conditions when this participant was excluded (active rest-TIS vs. active task-TIS: Fisher z = −2.17, p = .030).

## Discussion

The goal of the present study was to probe the functional role of the hippocampus in motor sequence learning and its consolidation at both the micro and macro timescales. To do so, we used Temporal Interference Stimulation (TIS) targeting the hippocampus of young healthy individuals in two separate experiments that differed based on the stimulation epochs, i.e., during task practice for the task-TIS experiment or during the short rest periods interspersed with task practice in the rest-TIS experiment. Results indicate that while hippocampal-targeted TIS did not modulate overall motor performance, rest-TIS enhanced consolidation at the micro timescale (i.e., between motor practice blocks). Although this effect did not carry over the 24h macro-consolidation interval at the group level, participants showing greater TIS-induced micro-offline gains in performance during initial learning showed greater overnight gains in performance at the macro timescale. Altogether, these results provide evidence that experimental interventions targeting the hippocampus modulate offline motor memory consolidation.

Our results indicate that hippocampal-targeted TIS delivered during inter-practice rest periods – but not control stimulation - modulated micro-online (i.e., within practice blocks) and micro-offline (i.e., during inter-practice rest intervals) changes in performance during initial motor sequence learning. Specifically, active TIS resulted in greater micro-offline gains and lower micro-online gains in performance relative to control stimulation, an effect that was not observed in the task-TIS experiment. These findings are in line with our a priori hypotheses based on prior work suggesting a critical role of the hippocampus in the micro-offline motor memory consolidation process (Buch et al., 2021; Chen et al., 2025; Gann et al., 2023; Jacobacci et al., 2020; Mylonas et al., 2024; Sjøgård et al., 2025) and further support the involvement of this structure in this process. We speculate that hippocampal-targeted TIS may have strengthened the reactivation of hippocampal task patterns during inter-practice rest periods, which in turn resulted in greater gains in performance across the micro rest intervals (Buch et al., 2021). The current results are also in line with two recent non-invasive brain stimulation studies using indirect modulation of hippocampal networks through prefrontal cortical transcranial magnetic stimulation (TMS) (Gann et al., 2023) or inhibitory transcranial ultrasound stimulation (TUS) of the hippocampus (Montenegro et al., 2025). Both studies showed that stimulation applied before learning hindered micro-offline gains in performance and shifted performance improvements toward the micro-online window. At the neural level, Gann et al. (2023) showed that stimulation not only resulted in a decrease in hippocampal activity during rest, but it also tended to reduce the persistence of hippocampal task patterns during the micro-rest intervals. While informative, these earlier studies report moderate effects of the stimulation interventions, and with the stimulation being applied prior to learning, it is impossible to disentangle the contribution of micro-online vs. micro-offline processes to the pattern of observed results. The present study is the first to successfully boost micro-offline consolidation through stimulation targeting the hippocampus during the specific rest time windows hosting the fast consolidation process.

Our findings also represent an important contribution to the ongoing debate regarding the mechanisms underlying micro-offline gains in performance. Non-memory-based frameworks have attributed these gains in performance to the dissipation of reactive inhibition (practice-induced performance slowing presumably related to fatigue buildup; Gupta & Rickard, 2022) or to the beneficial effect of motor preplanning (i.e., faster performance after a period of rest during which movement could presumably be pre-planned; Das et al., 2025) rather than to an active memory consolidation process. The present findings are difficult to fully reconcile with the dissipation-of-fatigue account, as we are unaware of any experimental evidence linking hippocampal function to reactive inhibition. However, evidence suggests that hippocampal patterns during motor preparation, i.e., before motor execution, predict the ordinal structure of upcoming learned motor sequences (Kornysheva et al., 2019; Yewbrey & Kornysheva, 2024). One could therefore argue that hippocampal-targeted TIS in the present study enhanced pre-planning processes and, in turn, increased the micro-offline gains in performance. The current dataset therefore does not allow us to conclude whether the hippocampal contribution is related to pre-or re-activation processes. In line with recent hybrid models suggesting that the amplitude of micro-offline gains in performance is likely the result of multiple – and perhaps interacting – factors (Brooks et al., 2026; Gupta & Rickard, 2024), it is possible that both pre-and re-activation of task patterns in the hippocampus contribute to greater micro-offline gains in performance.

Our results indicate no evidence for an effect of hippocampal-targeted TIS, regardless of whether it was applied during task practice or during inter-practice rest intervals, on changes in performance at the 24h retest. This was contrary to our expectations, as task-related hippocampal responses during initial MSL have been consistently linked to sleep-dependent motor memory consolidation in previous research (Albouy et al., 2008; Albouy, Fogel, et al., 2013; Albouy, King, et al., 2013). The null results observed in the task-TIS experiment might suggest that active task practice does not represent an optimal time window for strengthening hippocampal recruitment. This generally concurs with neuroimaging evidence showing negative BOLD signal in the hippocampus during MSL task practice (Albouy et al., 2015; Gann et al., 2023; Jacobacci et al., 2020). However, it is unclear why the beneficial effect of rest-TIS on consolidation at the micro timescale did not carry over the 24h retention interval. It is possible that TIS only had a transient effect on hippocampal plasticity during initial learning. This interpretation is consistent with the findings of Wessel et al. (2023), who showed that theta-burst patterned TIS targeting the striatum enhanced motor performance during training but that this effect did not persist at post-training, 90-minute, or 24-hour follow-up assessments. Similarly, albeit with a different type of approach, direct inhibitory TUS of the hippocampus did not produce group-level differences in overnight consolidation despite robust effects on micro timescale dynamics (Montenegro et al., 2025). Another possibility is that consolidation processes at the micro and macro timescales are independent and that interventions modulating the fast consolidation processes do not influence the slow consolidation process. Previous neuroimaging studies point against this view as similar hippocampal reactivation processes have been observed at both the micro and macro timescales (Gann et al., 2023; King et al., 2022) but prior behavioral studies reported mixed results with respect to this relationship. Specifically, some studies showed no evidence for a correlation between micro-and macro-offline changes in performance (e.g., Bönstrup et al., 2019; Jacobacci et al., 2020) while other work reported a positive association (e.g., Van Roy et al., 2024). The current results showed a significant positive correlation between these metrics but only in the active rest-TIS condition. These results suggest that although hippocampal-targeted TIS did not influence overnight macro changes in performance at the group level, participants with greater TIS-induced gains in performance at the micro timescale showed greater gains at the 24h retest. These results therefore suggest that the effect of hippocampal-targeted TIS carried over the 24h consolidation interval in participants who responded more strongly to the stimulation during initial learning. However, note that although this correlation was greater in the rest-TIS than in the task-TIS experiment, it did not differ between active and control conditions within the rest-TIS experiment, and it was strongly influenced by an extreme participant. The corresponding results should therefore be interpreted with caution.

An important limitation of the present study is the absence of individual structural MRI data, which prevented us from performing individualized head modeling to verify that the stimulation montage was effective in targeting the hippocampus in every participant. Instead, the electrode montage was adopted from previous studies that validated the procedure with cadaver electric field measurements, head modelling and fMRI data (Beanato et al., 2024; Violante et al., 2023). It is possible that individual anatomical and electrode positioning variability may have affected the focality of the stimulation and perhaps attenuated some of the group-level effects. In addition, although our exploratory electric field simulations showed that the targeted right hippocampus was strongly exposed (see Figure 1D-E), they also predicted that the right putamen - a brain region critically involved in MSL - received a field of comparable magnitude (see section 3 in the Supplemental information) which may have affected the present pattern of results. Based on previous research showing a preferential recruitment of the putamen during online task practice (Albouy, King, et al., 2013; Doyon et al., 2009; Van Roy et al., 2025) which can be strengthened by striatal-TIS (Wessel et al., 2023), one would expect the potential putamen stimulation in the current task-TIS experiment to enhance general motor performance during learning (Wessel et al., 2023). The data did not show such evidence. It is possible that simultaneous putamen-hippocampus stimulation during task practice might have interfered with the expected beneficial effects, but this remains speculative. In the same vein, as the putamen is known to not be activated during inter-practice rest intervals, it is unclear how putamen or simultaneous putamen-hippocampus stimulation might have influenced motor performance in the rest-TIS experiment. Future work should address these limitations by acquiring individual structural MRI data to enable individual electric field modeling prior to stimulation in order to tailor the electrode montage to optimize focality in each individual (Tang et al., 2025; Yang et al., 2024). In addition, combining TIS with concurrent functional neuroimaging would allow us to examine the effect of TIS on hippocampal (and striatal) activity and connectivity patterns during motor sequence learning and link stimulation-induced neural changes to the behavioral enhancements.

In conclusion, the present pre-registered study provides the first direct evidence that non-invasive stimulation targeting the hippocampus during the rest periods interspersed with motor practice enhances micro-offline motor memory consolidation, supporting a causal role of the hippocampus in offline motor memory consolidation processes. These findings challenge the long-standing dichotomization of memory systems and push the field to reconsider and further update models of memory system organization. They also open new avenues to optimize motor learning processes via the targeting of hippocampal-mediated processes, which may ultimately contribute to alleviating age- or pathology-related declines in motor learning.

## Methods

This study was preregistered in the Open Science Framework (OSF, available at https://osf.io/5m37g). The preregistration document detailed our design, hypotheses and the statistical models used to test our *a priori* hypotheses. Any analysis reported in this paper that was not preregistered is referred to as exploratory in the manuscript. There were no deviations from the preregistration with respect to the statistical plan but there were occasional issues that arose during data acquisition which resulted in deviations from the standard protocol (e.g., computer malfunction, experimental error etc.). These deviations are reported in Section 9 of the Supplements.

### Participants

Young healthy participants (mean age: 24.59 years ranging from 18 to 35) were recruited for this study and received monetary compensation for their time and effort. Written informed consent was obtained prior to participation. The study protocol was approved by the University of Utah Institutional Review Board (IRB_00183830) and conducted in accordance with the Declaration of Helsinki (World Medical Association, 2013). Inclusion criteria are: 1) right handed, as assessed by the Edinburgh Handedness Inventory (Oldfield, 1971), 2) no previous extensive training with a musical instrument or as a professional typist, 3) free of neurological, psychological, psychiatric, or sleep conditions, including depression and anxiety as assessed by the Beck’s Depression (Beck et al., 1996) and Anxiety (Beck et al., 1988) Inventories, 4) no indications of abnormal sleep, as assessed by the Pittsburgh Sleep Quality Index (Buysse et al., 1989) (PSQI); 5) not considered extreme morning or evening types, as quantified with the Horne & Ostberg chronotype questionnaire (Horne & Ostberg, 1976), 6) free of psychoactive or sleep-affecting medications; and 7) no contra-indications to brain stimulation procedures. None of the participants had a history of severe concussion that could affect brain structure, were shift workers, or had undertaken trans-meridian travel in the month preceding the study.

Sample size was determined through an *a priori* power analysis using GPower 3.1. The effect size estimate (Cohen’s d = 0.524) was derived from Violante et al. (2023), who used a similar 5 Hz hippocampal stimulation procedure and reported a ∼12% improvement in accuracy for declarative memory recall in the active versus control stimulation condition in a within-subject design. A two-tailed paired-sample t-test with α = 0.05 and power = 0.80 indicated that 31 participants would be required to detect this effect. We included 32 participants in each experiment (64 total) to allow for the counterbalancing between sequences, representations and stimulation type described in the procedure below.

Seventy-one participants were recruited for this study and were pseudo-randomly assigned (alternative assignment) to one of the two independent experiments (task-TIS experiment and rest-TIS experiment, see procedure). Out of the 36 participants who were assigned to the task-TIS experiment, one dropped out between sessions due to unexpected scheduling conflict, and three were excluded for non-compliance with sleep/wake instructions. Thirty-two participants (18 females, mean ± SD age: 25.09 ± 4.98 years) were eventually included in the analyses of the task-TIS experiment. Out of the 35 participants assigned to the rest-TIS experiment, one participant withdrew after the habituation phase due to discomfort related to the electrical stimulation, and two were excluded for not complying with the sleep/wake instructions. A total of 32 participants (18 females, mean ± SD age: 24.09 ± 4.13 years) were therefore included in the analyses of the rest-TIS experiment. Participant demographics, screening assessment scores, vigilance and sleep characteristics are presented in Supplemental section 1. Some of the protocol deviations described in Supplemental section 9 resulted in the exclusion of subsets of participants from specific statistical contrasts presented in this manuscript. The sample sizes for each analysis are explicitly stated in the corresponding figure or table captions.

### General Design

This study includes two experiments (task-TIS and rest-TIS) performed on independent participant samples. The two experiments are identical except for the epoch during which stimulation was delivered (i.e., during task practice for the task-TIS experiment and during inter-practice rest periods for the rest-TIS experiment). Within each experiment, participants completed two visits one week apart (referred to as Visit 1 in week 1 and Visit 2 in week 2), during which they received either active or control hippocampal-targeted TIS (see TIS section below) in a counterbalanced order (Figure 1A).

Each experimental visit included two consecutive testing days (Figure 1a). Participants were instructed to follow a regular sleep schedule (latest bedtime: 1am; minimum 7 hours of sleep) starting 3 days before Day 1 and between Day 1 and Day 2 of each experimental visit. Compliance to the sleep schedule was monitored using sleep diaries for the 3 days prior to Day 1 and wrist actigraphy (ActiGraph wGT3X-BT, Pensacola, FL) in combination with the sleep diary between experimental Day 1 and Day 2. All four experimental sessions (Days 1 and 2 of Visits 1 and 2) took place between 9am and 7pm and at approximately the same time of day (within a ±2-hour window) within each participant. At the beginning of each of these four experimental sessions, participants completed a brief questionnaire to collect information regarding their sleep patterns in the previous 24 hours (St. Mary’s Hospital Sleep Questionnaire; Ellis et al., 1981), their subjective feelings of alertness (Stanford Sleepiness Scale; Hoddes et al., 1973), and compliance with protocol restrictions (e.g., no caffeine, alcohol, strenuous exercise, tobacco, or psychoactive medication).

On Day 1, following the initial assessments mentioned above, the TIS electrodes were placed on the participants’ scalp (see TIS section below). Participants then performed a psychomotor vigilance task (PVT; Dinges & Powell, 1985) to obtain an objective measure of their alertness, followed by a pseudo-random serial reaction time task (SRTT; Nissen & Bullemer, 1987) to assess baseline motor performance. Next, participants were exposed to the electrical stimulation procedure with a stimulation habituation phase during which participant’s sensations were monitored and logged (see below). Participants were then familiarized with the motor sequence task with a short pre-training phase that was followed by two 20-block training phases (referred to as Training 1 and Training 2). Stimulation (active or control, depending on the visit) was delivered during practice blocks in the task-TIS experiment or during the inter-practice rest periods in the rest-TIS experiment. A sensation questionnaire assessing potential side effects resulting from the stimulation was administered after each training phase. Following Training 2, the stimulation electrodes were removed, and participants completed a short debriefing questionnaire to assess the effectiveness of blinding (see blinding effectiveness section below). Finally, a 4-block post-training test was administered without stimulation to obtain a stimulation-and fatigue-free measure of end-of-training performance (Cai & Rickard, 2009; Gupta & Rickard, 2022; Pan & Rickard, 2015). Participants were then sent home with an actiwatch and were asked to continue to respect the constant sleep schedule and comply with the restrictions mentioned above (e.g., no caffeine, alcohol, strenuous exercise or tobacco).

On Day 2, approximately 24 hours (i.e., ± 2 hours) after training, participants returned to the lab for a retest session. Upon arrival, participants completed the same questionnaires and compliance checks as on Day 1 and performed the PVT to assess alertness. Following these initial assessments, participants performed the motor task without stimulation. The session included a retest on the learned sequence (4 blocks) which was followed by two task representation tests to assess spatial (4 blocks) and motoric (4 blocks) representations of the learned task (order counterbalanced across participants). Last, participants were tested on a novel sequence (4 blocks). The Day 2 session of each visit concluded with a short debriefing questionnaire on the motor task.

### Tasks

All tasks were implemented in MATLAB using Psychtoolbox version 3 (Brainard, 1997; Kleiner et al., 2007).

### Psychomotor Vigilance Test (PVT)

Participants performed a psychomotor vigilance task (PVT; Dinges & Powell, 1985) to obtain an objective measure of their alertness. During the PVT, participants were asked to fixate on a cross in the center of the screen and to press the space bar as fast as possible when the fixation cross disappeared. A timer appeared on the screen displaying the participants’ reaction time in milliseconds. Participants completed 35 trials of the PVT task, and vigilance was indexed as the median reaction time across all trials.

### Serial Reaction Time Task (SRTT)

A random version of the SRTT was used to ensure that baseline motor performance was equivalent between experimental visits (within participants) and experiments (between participants). During this task (Nissen & Bullemer, 1987), four squares were presented on the screen, each corresponding to one of the four keys on a specialized keyboard and to one of the four fingers (no thumbs) of the non-dominant left hand. The color of the outline of the squares alternated between red and green, indicating rest and practice blocks, respectively. During the practice blocks, participants pressed as quickly as possible the key corresponding to the location of a green filled square that appeared on the screen. After a response, the next square in the stream immediately changed to green following a pseudo-random order generated by concatenating 16 randomized permutations of the four possible positions (64 key presses per block). Participants completed 4 practice blocks that were separated by 15-s rest intervals. Speed (mean reaction time on correct trials, from cue onset to key press) and accuracy (percentage of correct responses) were used as outcome measures.

### Motor Sequence Learning Task (MSL)

Participants performed a unimanual sequential finger-tapping task similar to our previous research (e.g., Albouy et al., 2015; Albouy, Fogel, et al., 2013). The task required participants to tap an eight-element finger sequence (4 fingers, no thumb) on a specialized keyboard as rapidly and accurately as possible with their left non-dominant hand. Motor performance on this task was assessed in terms of speed (mean inter-response interval [IRI] on correct keypresses) and accuracy (percentage of correct transitions within a block), which served as the main outcome variables. The sequences to perform were explicitly provided to the participants and continuously displayed on the screen during task practice. On Day 1, for each visit (weeks 1 and 2), the MSL session started with a brief pre-training phase during which participants performed the sequence repeatedly and slowly until three consecutive correct sequences were completed. The training session then consisted of 2 training phases of 20 practice blocks each (Training 1 and 2) and was followed by an immediate, stimulation-free post-training test (after a ∼2-min break) of 4 practice blocks in order to minimize the confounding effect of fatigue on end-training performance (Cai & Rickard, 2009; Gupta & Rickard, 2022; Pan & Rickard, 2015). Different sequences were learned in the two experimental visits one week apart (order counterbalanced across participants). The learned sequences were 41321342 and 23142143 (Sets A and B, respectively), whereby 1 and 4 represent the index and the little finger, respectively. On Day 2 of each experimental visit, participants were retested on the sequence learned on Day 1 with 4 blocks of practice. In addition, to probe the nature of the learned representations, participants performed two representation tests (order counterbalanced across participants) with an inverted keyboard setup (Figure 1A) and using a procedure similar to our prior research that affords the assessment of the spatial and motoric representations of the learned sequence (Albouy et al., 2015; Albouy, Fogel, et al., 2013). To assess the spatial representation, participants performed a mirrored version of the sequence learned on Day 1 using the inverted keyboard (i.e., spatial sequence A: 14234213 and spatial sequence B: 32413412, Figure 1A). As a result, the required finger movements were different compared to the learned sequence, but the presses occurred at the same spatial locations as during training. To assess the motor representation, participants performed the same sequence of movements as during training (i.e., motor sequence A: 41321342, motor sequence B: 23142143, see Figure 1A), which preserved the motoric representation of the task but resulted in a different spatial representation. Last, the keyboard and hand were flipped again to the standard position and participants performed a new sequence (new sequences A: 32141243 and B: 13242431) to test for sequence-specificity effects. All eight possible combinations of the sequence order (A-B/B-A), stimulation order (active-control/control-active), and representation-test order (Spatial-Motor/Motor-Spatial) were balanced in each experiment (4 repetitions of each of the 8 combinations in each experiment of 32 participants).

For all the task conditions mentioned above, the practice blocks were indicated with a green square displayed in the middle of the screen with the sequence of numbers shown slightly above. Each practice block lasted 30 s after which the sequence of numbers disappeared and the central square automatically turned red, indicating a rest period (duration 30 s). During the practice blocks, participants were instructed to continuously tap the sequence in a self-initiated manner until the stop signal (red square) was given. During rest periods, participants were instructed to keep their fingers still and look at the red square. TIS was applied during Training phases 1 and 2 only (i.e., not during Test and Retest sessions) and was administered during practice blocks for the task-TIS experiment and during the inter-practice rest periods in the rest-TIS experiment.

### Transcranial Temporal Interference Stimulation (TIS)

TIS was delivered using the LOOP-IT closed-loop neuromodulation system (neuroConn GmbH, Ilmenau, Germany) controlled via a customed MATLAB script (R2023b; MathWorks, Natick, MA, USA) with Psychtoolbox-3 and a National Instruments USB-6341 DAQ device.

The electrode montage was adapted from Violante et al. (2023) and Beanato et al. (2024) to target the right hippocampus. The right hippocampus was selected as the stimulation target based on neuroimaging evidence that hippocampal recruitment during MSL supports the encoding of the spatio-temporal representation of the motor sequence, i.e., the ordinal arrangement of movements in space and time (Albouy et al., 2015; Dolfen et al., 2024). The montage included two pairs of electrodes placed on FT8, Fp1, TP8 and P7 electrode positions on the EEG 10-10 system. Following skin preparation with alcohol, 3 cm² circular conductive rubber electrodes were coated with an approximately 1-mm layer of conductive paste (Ten20, Weaver and Company, or Abralyt HiCl, Easycap GmbH) and affixed to the marked positions. Electrodes were secured with medical tape (3M micropore tape) and a tubular elastic retainer net to maintain contact. Impedances were measured with an electrode impedance checker (EL-CHECK, BIOPAC Systems Inc., Goleta, CA, USA; test frequency: 25 Hz; test current: 3.5 µA RMS), monitored throughout the experiment and maintained below 20 kΩ (von Conta et al., 2022; Wessel et al., 2023). In the task-TIS experiment, mean impedance values were comparable between the active (M = 14.09 kΩ, SD = 4.73) and control (M = 15.37 kΩ, SD = 5.57) stimulation conditions (t(31) = −1.76, p = .089) throughout the experiment. Similarly, in the rest-TIS experiment, impedance did not differ between the active (M = 13.52 kΩ, SD = 5.38) and control (M = 15.82 kΩ, SD = 8.15) stimulation conditions (t(31) = −1.50, p = .144) throughout the experiment.

In each experiment, two stimulation conditions were employed in a within-subject design (7-9 days washout period between the two stimulation sessions). For *active* stimulation, two sinusoidal currents of 2000 Hz and 2005 Hz were delivered through electrodes pairs TP8-P7 and FT8-Fp1 at 2 mA (baseline-to-peak) per channel, producing a 5 Hz envelope targeting the right hippocampus. For *control* stimulation, two identical 2000 Hz currents were applied using the same electrode configuration, producing a flat envelope without amplitude modulation (Beanato et al., 2024; Wessel et al., 2023). Prior to the main stimulation session, participants completed a habituation stimulation session (HAB in Figure 1A) during which they were exposed to 4 blocks of 30 s of stimulation (including 5 s ramp-up and -down) at four stepwise current intensities (0.5, 1, 1.5 and 2 mA). Stimulation was then delivered during initial motor sequence learning in blocks of 30 s, including a 5 s ramp-up and a 5 s ramp-down phase. In the task-TIS experiment, stimulation was delivered during each 30-s practice block of the two training sessions (i.e., 40 practice blocks stimulated in total, stimulation duration = 20 min for each [active / control] experimental visit). In the rest-TIS experiment, stimulation was delivered during each 30-s rest block following a practice block during the two training sessions (i.e., 40 rest blocks stimulated in total, stimulation duration = 20 min for each [active / control] experimental visit).

Sensations associated with the stimulation were assessed with two questionnaires administered during the habituation session and after each training phase. During the habituation stimulation phase, participants reported any perceived sensations after each block of stimulation and, if present, rated their intensity on a scale from 1 (mild) to 3 (strong) and provided adjectives best describing the sensation. In addition, after each stimulation session (i.e., after Training 1 and Training 2), participants completed a sensation questionnaire adapted from Brunoni et al. (2011) and consistent with previous TIS studies (Violante et al., 2023; Wessel et al., 2023). Participants were asked to rate the intensity of 15 sensations (itching, scalp pain, burning, warmth, metallic taste, fatigue, headache, tingling, sleepiness, mood change, visual sensation, dizziness/nausea, nervousness/anxiety, discomfort and other sensations) on a scale from 0 (absent) to 3 (severe). Last, blinding effectiveness was assessed using a debriefing questionnaire administered at the end of each experimental visit, adapted from Violante et al. (2023) and Greinacher et al. (2019). Participants were asked whether they believed they received the active stimulation condition during the session (yes/no) and to rate the confidence of their response on a scale from 1 (not confident at all) to 10 (extremely confident). A weighted score was computed by multiplying the response (yes = +1; no = −1) by the confidence rating, such that higher positive scores indicate greater confidence in having received active stimulation. Participants who perceived the stimulation were also asked whether they believed the stimulation improved their performance, disrupted their performance, or had no particular effect.

### Behavioral data analyses

#### Preprocessing and outcome measures

Consistent with our preregistration, primary analyses focused on performance speed and results related to performance accuracy are reported in Supplemental section 5. Individual IRIs were excluded from the analyses if they fell more than 3 standard deviations above or below the participant’s mean IRI for that block. The total number of discarded trials (inaccurate and outlier) was 5.3% across training sessions and conditions (5.4% and 5.4% in active and control visits, respectively in the task-TIS experiment; 5.2% and 5.4% in active and control visits in the rest-TIS experiment, respectively; see Supplemental section 10 for detailed statistical analyses).

Micro-scale learning was quantified by computing within-block (micro-online) and between-block (micro-offline) changes in performance. Micro-online changes in performance were calculated as the difference in mean correct IRI (cIRI) between the first eight and the last eight correct transitions within each block, then summed across blocks. Micro-offline changes in performance were calculated as the difference in mean cIRI between the last eight correct transitions of one block and the first eight correct transitions of the subsequent block, then summed across the 40 training blocks. In addition to these pre-registered analyses, we performed exploratory analyses to examine the amplitude of these micro-offline gains in performance in an earlier learning time-window. Micro-offline gains in performance have indeed been described to be most prominent during the early phase of learning, when the learning curve is steep and within-block performance decrements are minimal (Bönstrup et al., 2019, 2020; Brooks et al., 2026). Earlier work examined micro-offline gains in performance across blocks accounting for 95% of the total initial learning (Bönstrup et al., 2019, 2020). In our data set, 86% of the total learning was reached at the end of Training 1 (with the total learning defined as the difference in group-mean cIRI between the first training block and the mean of the last four training blocks (blocks 37–40)). We therefore performed exploratory analyses on this earlier learning stage for which micro-online and micro-offline changes in performance were summed over Training 1 only (i.e., the first 20 practice blocks and the 19 intervening rest periods). The corresponding results are reported in Supplemental section 7 and are similar to those observed using both Training 1 and Training 2 data presented in the main text.

Macro-offline changes in performance were computed as the difference in mean cIRI between the four test blocks at the end of Day 1 training and the four retest blocks on the learned sequence on Day 2.

Transfer indices were computed separately for the spatial and motor representations of the trained sequence. For each representation type, the transfer index was calculated as the difference between the mean cIRI across the four blocks of the novel sequence task and the mean cIRI across the four blocks of the spatial/motor representation task such that positive values indicate faster performance on the representation task relative to the novel sequence.

#### Statistical analyses

Statistical analyses were performed using R (R Core Team, 2023). Statistical tests were considered significant at p < 0.05. For repeated-measures ANOVAs, Greenhouse-Geisser corrections were applied when the assumption of sphericity was violated. When necessary, corrections for multiple comparisons were conducted using the False Discovery Rate (FDR) procedure within each family of hypothesis tests. Effect sizes are reported using Cohen’s dz (i.e., Cohen’s d for paired-samples designs, computed as the mean difference divided by the standard deviation of the differences) for paired-samples t-tests and partial η² for repeated-measures ANOVAs.

In our confirmatory analyses, we first tested whether performance during training differed between stimulation conditions within each experiment. To do so, 2 (Training phase: Training 1 vs. Training 2) × 20 (Block) × 2 (Stimulation condition: Active vs. Control) repeated-measures ANOVAs were performed separately for each experiment. For the post-training test, 4 (Block) × 2 (Stimulation condition) repeated-measures ANOVAs were conducted. These analyses were also performed with the additional between-subject factor Experiment (2: task-TIS vs. rest-TIS, results reported in Supplemental section 4).

Second, we tested whether micro-online and micro-offline changes in performance differed between stimulation conditions using 2 (Micro-gain: online vs. offline) × 2 (Stimulation condition: Active vs. Control) repeated-measures ANOVAs performed on the sum of micro-online and -offline changes in performance across the 40 training blocks for each experiment. These analyses were then repeated with an additional between-subject factor Experiment (2, results reported in Supplemental section 7). In exploratory analyses focusing on an earlier learning phase, the same 2 (Micro-gain) × 2 (Stimulation condition) ANOVAs and follow-up paired t-tests were performed on micro-online and micro-offline changes in performance summed over Training 1 only (Supplemental section 7).

Third, we tested whether the macro-offline changes in performance differed between stimulation conditions using paired t-tests within each experiment, and subsequently with a 2 x 2 ANOVA using stimulation condition as within-subject factor and Experiment as between-subject factor (results reported in Supplemental section 8).

Fourth, Pearson correlations between micro-offline and macro-offline changes were computed and compared between stimulation conditions within each experiment, and between experiments.

Fifth, we tested whether stimulation affected spatial and motor representations of the learned sequence. To do so, 2 (Representation: motor vs. spatial) × 4 (Block) × 2 (Stimulation condition: Active vs. Control) repeated-measures ANOVAs were performed for each experiment. Additionally, 2 (Transfer index: motor vs. spatial) × 2 (Stimulation condition) repeated-measures ANOVAs were conducted. These analyses were repeated using the additional between-subject factor Experiment (2). Corresponding results are reported in Supplemental section 6).

Control analyses were performed to verify that the pattern of results emerged from our experimental manipulation rather than from potential confounding factors. These included comparisons of baseline motor performance (SRTT) between stimulation conditions as well as assessment of potential sequence type (Set A vs. Set B) and order effects (Visit 1 vs. Visit 2). Results corresponding to these control analyses are provided in Supplemental section 11. Overall, these control analyses confirmed that baseline motor performance did not differ between stimulation conditions in either experiment. Results also show that performance on one sequence set was faster than the other set in the task-TIS experiment and that performance during Visit 2 was better than during Visit 1 during initial training in both experiments. However, as sequence sets and stimulation order were fully counterbalanced across conditions and experiments, it is unlikely that these factors confounded the reported results.

### Exploratory analyses on sensation and blinding

#### Sensation analyses

Sensation ratings were ordinal, non-normally distributed, and occasionally missing. They were therefore analyzed with cumulative link mixed models (CLMMs; clmm function, ordinal package; Christensen, 2023), which model the ordered rating categories directly rather than assuming an interval scale, and accommodate unbalanced data. Analyses of sensation ratings collected during the habituation session were performed within each experiment using stimulation condition (2: active vs. control) and current intensity (4: 0.5, 1.0, 1.5, 2.0 mA) as fixed factors, and random intercepts for participant and for participant-by-stimulation-condition to account for potential rating fluctuations between the separate experimental visits (each stimulation condition being administered in a separate visit). Between-experiment comparisons were conducted using the same model with experiment (2: task-TIS vs. rest-TIS) as an additional fixed factor. Statistical significance of the fixed effects was determined using Type II likelihood-ratio χ² tests, and post hoc pairwise comparisons were performed using estimated marginal means with Tukey adjustment (emmeans package; Lenth, 2024).

For the post-training sensation questionnaires, the 15 sensation items were distributed in eight categories similar as in previous research (Wessel et al., 2023) and consisting of itching, pain (including headache and scalp pain), burning, warmth/heat, tingling, metallic/iron taste, fatigue/decreased alertness (including sleepiness), and other (including mood change, visual sensation, dizziness/nausea, nervousness/anxiety, discomfort, and other sensation). For categories comprising multiple items, the maximum rating across the constituent items was used as the category score. Category scores from the two training phases of each visit were analyzed within each experiment using CLMMs with stimulation condition (2: active vs. control), sensation category (8) and training phase (2: phase 1 vs. phase 2) as fixed factors, and random intercepts for participant and for stimulation condition nested within participant (participant-by-stimulation-condition). Between-experiment comparisons included experiment (2: task-TIS vs. rest-TIS) as an additional fixed factor. The same inference procedure (Type II likelihood-ratio χ² tests, Tukey-adjusted post hoc comparisons, and linear mixed model robustness checks) was applied as described above (Supplemental section 2).

#### Blinding analyses

As the weighted condition-identification scores are ordinal in nature, they were analyzed with CLMMs as described above. Within each experiment, weighted scores were compared between stimulation conditions using a CLMM with stimulation condition (2: active vs. control) as a fixed factor and a random intercept for participant. A participant-by-stimulation-condition intercept was not included here because only one weighted score was obtained per participant and visit. Between-experiment comparisons were conducted using the same model with experiment (2: task-TIS vs. rest-TIS) as an additional fixed factor. Statistical significance of the fixed effects was determined using Type II likelihood-ratio χ² tests. In addition, weighted scores were tested against zero (i.e., chance-level identification) within each experiment and stimulation condition using Wilcoxon signed-rank tests. The distribution of responses on the perceived effect on performance was compared between conditions within each experiment using Fisher’s exact tests and between experiments using Cochran–Mantel– Haenszel tests (Supplemental section 2).

### Exploratory electric field modeling

We performed simulation analyses to estimate the exposure of the target region to the low-frequency envelope modulation of the field. The electric fields induced by the experimental montage were simulated on the MIDA head model (Iacono et al., 2015), a high-resolution (0.5 mm) anatomical model of the head and neck of a healthy 29-year-old female volunteer that distinguishes 153 anatomical structures (Iacono et al., 2015) and served as reference head model in the study that validated the present montage (Violante et al., 2023). The Temporal Interference Planning (TIP; TIP.lite v5.2.3; <u>IT’IS Foundation, 2024</u>) model used here includes 115 distinguished tissues and regions with separate scalp layers, cortical and cancellous bone, dura, cerebrospinal fluid, grey and white matter and the deep brain nuclei. Anisotropic white matter and midbrain conductivity was derived from the diffusion tensor imaging data provided with the model in the TIP toolbox. All remaining tissue conductivity values were assigned according to the low-frequency conductivity section of the IT’IS Tissue Properties Database (version 4.1; <u>Hasgall et al., 2022)</u>. The model was exported to Sim4Life.lite (v9.6; ZMT Zurich MedTech AG, Zurich, Switzerland), in which the two electrode pairs were simulated separately. Montage and electrode parameters matched the experiments: 3 cm² circular electrodes (radius 9.8 mm) were placed on the scalp at the Fp1, FT8, TP8 and P7 positions of the EEG 10-10 system. The electric field was computed with the ohmic-current-dominated electro-quasistatic solver on a rectilinear grid with a maximum cell size of 1 mm within the brain and skull (0.75 mm in the electrodes, 2 mm in the remaining head and neck), resulting in about 10 million voxels per simulation. Dirichlet voltage boundary conditions were applied to the electrodes of each pair, followed by current normalization to 2 mA per channel, and the electrode–scalp contact was treated as ideal. The TIS electric field was quantified voxel-wise as the maximal amplitude of the low-frequency envelope modulation of the superimposed fields (TI_max; <u>Grossman et al., 2017)</u> after linear resampling of both fields onto a common 1 mm isotropic grid, and is displayed in grey matter in the MNI space (Figure 1D). For the region-of-interest analysis (Figure 1E), the left and right hippocampi were defined from the hippocampus label of the MIDA tissue segmentation, which covers both hemispheres, split into left and right at the midsagittal plane of the model, and the field was extracted from all voxels within each mask. Control regions were defined in the same way for the putamen, caudate and thalamus (derived from the MIDA segmentation). For the primary motor cortex M1, grey matter volumes within two 10-mm-radius spheres centered on the left and right hand areas of M1 (MNI x, y, z = ±36, −22, 57 mm) were transferred to the MIDA model by nonlinear registration of the MIDA T1-weighted image to the MNI152 ICBM 2009a template (Fonov et al., 2011) using DIPY v1.12 (affine followed by symmetric diffeomorphic registration; <u>Garyfallidis et al., 2014)</u>. The corresponding exposure distributions are shown in Supplemental section 3.

## Data availability

The datasets generated and analyzed, along with the analysis scripts used in the current study, will be made available in a public data repository upon publication.

## Acknowledgments

This work provided pilot data that supported the development of a National Institute of Mental Health of the National Institutes of Health funded project under award number R21MH140335 (to G.A.). Additional support was provided by University of Utah internal funds and by equipment available in the College of Health Research Facility. We thank Sophia Terrill and Elise Perry for their assistance with data collection, and all participants for their time and commitment. I.R.V is supported by the Biotechnology and Biological Sciences Research Council (BB/Y011856/1).

## Author contributions

Conceptualization and funding acquisitions: GA, BRK, CI and NG; Methodology: XP, GA, BRK, IRV, CI and NG; Investigation: XP and JT; Formal analyses: XP and GA; Data curation and visualization: XP; Writing – original draft: XP and GA; Writing – review & editing: XP, JT, IRV, NG, BRK and GA. Project administration: GA and BRK. Supervision: GA.

