## Supplemental Information for "Non-invasive temporal interference stimulation targeting the hippocampus enhances fast motor memory consolidation"

### 1. Participants demographics, sleep and vigilance characteristics

**Table S1. Participants' demographics.**

| Measure / Effect | task-TIS (N = 32) | rest-TIS (N = 32) | Between experiments |
|---|---|---|---|
| Sex (female / male, n) | 18 / 14 | 18 / 14 | $\chi^2(1)$ = 0.00, p = 1.000 |
| Age (years) | 25.1 ± 5.0 | 24.1 ± 4.1 | t(59.9) = 0.87, p = .385 |
| Edinburgh Handedness | 84.7 ± 18.4 | 86.1 ± 14.5 | t(58.7) = −0.34, p = .736 |
| BDI score | 3.4 ± 4.8 | 3.5 ± 3.9 | t(59.3) = −0.06, p = .955 |
| BAI score | 3.1 ± 4.3 | 2.8 ± 2.7 | t(51.8) = 0.38, p = .703 |
| PSQI score | 3.1 ± 1.8 | 3.8 ± 2.4 | t(57.9) = −1.26, p = .214 |
| Chronotype | 50.6 ± 9.6 | 52.7 ± 9.6 | t(62.0) = −0.87, p = .385 |
| Epworth Sleepiness Scale | 4.9 ± 4.1 | 5.4 ± 4.4 | t(61.6) = −0.47, p = .639 |

*Note.* Values are mean ± SD. Demographics were compared between experiments using Welch's independent-samples t-tests (continuous variables) or a $\chi^2$ test (sex). BDI = Beck Depression Inventory; BAI = Beck Anxiety Inventory; PSQI = Pittsburgh Sleep Quality Index; Chronotype = Horne & Östberg Morningness–Eveningness Questionnaire.

**Table S2. Sleep and vigilance characteristics.**

| | | |
|---|---|---|
| **Sleep duration (min)[a]** | | |
| ***Active stimulation*** | | |
| Night 1 | 449.7 ± 179.8 | 446.4 ± 155.6 |
| Night 2 | 463.0 ± 135.4 | 457.6 ± 159.0 |
| Night 3 | 458.1 ± 127.3 | 434.5 ± 174.5 |
| Night 4[b] | 478.1 ± 46.3 | 478.3 ± 51.0 |
| RM ANOVA: Night (4) | F(2.04, 63.07) = 0.48, p = .627, $\eta^2$p = 0.015 | F(1.78, 55.25) = 1.18, p = .311, $\eta^2$p = 0.037 |
| ***Control stimulation*** | | |
| Night 1 | 460.4 ± 161.2 | 470.2 ± 175.1 |
| Night 2 | 444.9 ± 156.6 | 454.1 ± 162.1 |
| Night 3 | 442.8 ± 156.6 | 454.9 ± 153.9 |
| Night 4[b] | 474.2 ± 47.7 | 467.3 ± 40.2 |
| RM ANOVA: Night (4) | F(1.66, 51.45) = 0.96, p = .376, $\eta^2$p = 0.030 | F(1.43, 44.42) = 0.24, p = .714, $\eta^2$p = 0.008 |
| RM ANOVA: Stimulation condition (2) × Night (4) | Stimulation condition, F(1.00, 31.00) = 0.21, p = .651, $\eta^2$p = 0.007; Night, F(1.74, 53.95) = 0.72, p = .472, $\eta^2$p = 0.023; Stimulation condition × Night, F(2.17, 67.44) = 0.59, p = .573, $\eta^2$p = 0.019 | Stimulation condition, F(1.00, 31.00) = 0.14, p = .712, $\eta^2$p = 0.004; Night, F(1.43, 44.37) = 0.62, p = .492, $\eta^2$p = 0.020; Stimulation condition × Night, F(2.16, 66.84) = 0.97, p = .388, $\eta^2$p = 0.031 |
| Mixed ANOVA: Experiment (2) × Stimulation condition (2) × Night (4) (N = 64) | Experiment, F(1.00, 62.00) = 0.00, p = .969, $\eta^2$p = 0.000; Experiment × Stimulation condition, F(1.00, 62.00) = 0.33, p = .570, $\eta^2$p = 0.005; Experiment × Night, F(1.59, 98.68) = 0.04, p = .925, $\eta^2$p = 0.001; Experiment × Stimulation condition × Night, F(2.51, 155.80) = 0.51, p = .642, $\eta^2$p = 0.008 | |
| **St. Mary's Hospital Sleep Questionnaire — Quality** | | |
| ***Active stimulation*** | | |
| Night 3 | 4.16 ± 0.77 | 4.13 ± 0.81 (n = 31) |
| Night 4[b] | 4.19 ± 0.54 | 4.16 ± 0.64 (n = 31) |
| RM ANOVA: Night (2) | F(1.00, 31.00) = 0.04, p = .851, $\eta^2$p = 0.001 | F(1.00, 30.00) = 0.06, p = .813, $\eta^2$p = 0.002 |
| ***Control stimulation*** | | |
| Night 3 | 4.00 ± 0.76 | 4.03 ± 0.74 |
| Night 4[b] | 4.25 ± 0.51 | 3.97 ± 0.86 |
| RM ANOVA: Night (2) | **F(1.00, 31.00) = 4.43, p = .044, $\eta^2$p = 0.125** | F(1.00, 31.00) = 0.16, p = .690, $\eta^2$p = 0.005 |

| | | |
|---|---|---|
| RM ANOVA: Stimulation condition (2) × Night (2); rest-TIS n = 31 | Stimulation condition, F(1.00, 31.00) = 0.18, p = .675, η²p = 0.006; Night, F(1.00, 31.00) = 2.04, p = .163, η²p = 0.062; Stimulation condition × Night, F(1.00, 31.00) = 1.09, p = .304, η²p = 0.034 | Stimulation condition, F(1.00, 30.00) = 2.50, p = .125, η²p = 0.077; Night, F(1.00, 30.00) = 0.15, p = .702, η²p = 0.005; Stimulation condition × Night, F(1.00, 30.00) = 0.29, p = .595, η²p = 0.010 |
| Mixed ANOVA: Experiment (2) × Stimulation condition (2) × Night (2) (N = 63) | Experiment, F(1.00, 61.00) = 0.44, p = .509, η²p = 0.007; Experiment × Stimulation condition, F(1.00, 61.00) = 0.57, p = .451, η²p = 0.009; Experiment × Night, F(1.00, 61.00) = 1.78, p = .187, η²p = 0.028; Experiment × Stimulation condition × Night, F(1.00, 61.00) = 1.20, p = .278, η²p = 0.019 | |
| **St. Mary's Hospital Sleep Questionnaire — Duration (min)** | | |
| ***Active stimulation*** | | |
| Night 3 | 480.2 ± 53.5 (n = 31) | 471.8 ± 41.4 (n = 31) |
| Night 4[b] | 479.3 ± 46.7 (n = 31) | 480.9 ± 52.9 (n = 31) |
| RM ANOVA: Night (2) | F(1.00, 30.00) = 0.01, p = .918, η²p = 0.000 | F(1.00, 30.00) = 1.33, p = .258, η²p = 0.042 |
| ***Control stimulation*** | | |
| Night 3 | 477.4 ± 61.7 (n = 31) | 502.6 ± 134.2 |
| Night 4[b] | 482.4 ± 64.5 (n = 31) | 471.7 ± 41.2 |
| RM ANOVA: Night (2) | F(1.00, 30.00) = 0.15, p = .698, η²p = 0.005 | F(1.00, 31.00) = 1.94, p = .174, η²p = 0.059 |
| RM ANOVA: Stimulation condition (2) × Night (2); task-TIS n = 30, rest-TIS n = 31 | Stimulation condition, F(1.00, 29.00) = 0.00, p = .987, η²p = 0.000; Night, F(1.00, 29.00) = 0.11, p = .741, η²p = 0.004; Stimulation condition × Night, F(1.00, 29.00) = 0.14, p = .707, η²p = 0.005 | Stimulation condition, F(1.00, 30.00) = 0.82, p = .374, η²p = 0.026; Night, F(1.00, 30.00) = 0.80, p = .378, η²p = 0.026; Stimulation condition × Night, F(1.00, 30.00) = 3.20, p = .084, η²p = 0.096 |
| Mixed ANOVA: Experiment (2) × Stimulation condition (2) × Night (2) (N = 61) | Experiment, F(1.00, 59.00) = 0.04, p = .842, η²p = 0.001; Experiment × Stimulation condition, F(1.00, 59.00) = 0.63, p = .432, η²p = 0.011; Experiment × Night, F(1.00, 59.00) = 0.86, p = .356, η²p = 0.014; Experiment × Stimulation condition × Night, F(1.00, 59.00) = 2.76, p = .102, η²p = 0.045 | |
| **Stanford Sleepiness Scale (SSS)** | | |
| ***Active stimulation*** | | |
| Day 1 | 2.06 ± 0.67 | 2.10 ± 1.04 (n = 31) |
| Day 2 | 2.03 ± 0.65 | 1.84 ± 0.78 (n = 31) |
| RM ANOVA: Day (2) | F(1.00, 31.00) = 0.09, p = .768, η²p = 0.003 | **F(1.00, 30.00) = 5.19, p = .030, η²p = 0.147** |
| ***Control stimulation*** | | |
| Day 1 | 2.19 ± 0.82 | 2.16 ± 0.92 |
| Day 2 | 1.91 ± 0.82 | 2.12 ± 1.01 |
| RM ANOVA: Day (2) | F(1.00, 31.00) = 2.76, p = .107, η²p = 0.082 | F(1.00, 31.00) = 0.05, p = .831, η²p = 0.001 |
| RM ANOVA: Stimulation condition (2) × Day (2); rest-TIS n = 31 | Stimulation condition, F(1.00, 31.00) = 0.00, p = 1.000, η²p = 0.000; Day, F(1.00, 31.00) = 2.95, p = .096, η²p = 0.087; Stimulation condition × Day, F(1.00, 31.00) = 1.35, p = .255, η²p = 0.042 | Stimulation condition, F(1.00, 30.00) = 3.34, p = .078, η²p = 0.100; Day, F(1.00, 30.00) = 2.15, p = .153, η²p = 0.067; Stimulation condition × Day, F(1.00, 30.00) = 1.61, p = .214, η²p = 0.051 |
| Mixed ANOVA: Experiment (2) × Stimulation condition (2) × Day (2) (N = 63) | Experiment, F(1.00, 61.00) = 0.00, p = .957, η²p = 0.000; Experiment × Stimulation condition, F(1.00, 61.00) = 1.46, p = .232, η²p = 0.023; Experiment × Day, F(1.00, 61.00) = 0.01, p = .934, η²p = 0.000; Experiment × Stimulation condition × Day, F(1.00, 61.00) = 2.88, p = .095, η²p = 0.045 | |
| **Psychomotor Vigilance Task (PVT, ms)[c]** | | |
| ***Active stimulation*** | | |

| | | |
|---|---|---|
| Day 1 | 308.0 ± 29.5 | 303.9 ± 31.6 |
| Day 2 | 308.1 ± 32.4 | 303.0 ± 32.4 |
| RM ANOVA: Day (2) | $F(1.00, 31.00) = 0.00$, $p = .988$, $\eta^2p = 0.000$ | $F(1.00, 31.00) = 0.10$, $p = .758$, $\eta^2p = 0.003$ |
| ***Control stimulation*** | | |
| Day 1 | 303.3 ± 24.7 | 308.1 ± 29.3 |
| Day 2 | 302.0 ± 30.5 | 306.8 ± 29.3 |
| RM ANOVA: Day (2) | $F(1.00, 31.00) = 0.19$, $p = .668$, $\eta^2p = 0.006$ | $F(1.00, 31.00) = 0.25$, $p = .621$, $\eta^2p = 0.008$ |
| RM ANOVA: Stimulation condition (2) × Day (2) | Stimulation condition, $F(1.00, 31.00) = 2.75$, $p = .107$, $\eta^2p = 0.081$; Day, $F(1.00, 31.00) = 0.14$, $p = .712$, $\eta^2p = 0.004$; Stimulation condition × Day, $F(1.00, 31.00) = 0.07$, $p = .800$, $\eta^2p = 0.002$ | Stimulation condition, $F(1.00, 31.00) = 1.18$, $p = .285$, $\eta^2p = 0.037$; Day, $F(1.00, 31.00) = 0.29$, $p = .592$, $\eta^2p = 0.009$; Stimulation condition × Day, $F(1.00, 31.00) = 0.01$, $p = .922$, $\eta^2p = 0.000$ |
| Mixed ANOVA: Experiment (2) × Stimulation condition (2) × Day (2) (N = 64) | Experiment, $F(1.00, 62.00) = 0.00$, $p = .987$, $\eta^2p = 0.000$; Experiment × Stimulation condition, $F(1.00, 62.00) = 3.67$, $p = .060$, $\eta^2p = 0.056$; Experiment × Day, $F(1.00, 62.00) = 0.03$, $p = .860$, $\eta^2p = 0.001$; Experiment × Stimulation condition × Day, $F(1.00, 62.00) = 0.02$, $p = .879$, $\eta^2p = 0.000$ | |

*Note.* Values are mean ± SD. RM ANOVA = repeated-measures ANOVA within each experiment (Greenhouse–Geisser correction applied to within-subject factors with more than one numerator df). The mixed ANOVA adds Experiment (task-TIS vs. rest-TIS) as a between-subject factor. Where noted, $n < 32$ due to missing data; analyses including these measures were run on the reduced samples. Bold $p$ values indicate significant effects ($p < .05$). [a] Sleep duration for Nights 1–3 was derived from sleep diaries; Night 4 sleep duration was computed as the mean of actigraphy and St. Mary's Hospital Sleep Questionnaire data (where actigraphy data were missing, only St. Mary's questionnaire data were used). [b] Night 4 = night between Day 1 and Day 2. [c] Median reaction time (ms) computed across trials of each stimulation condition.

### 2. Exploratory analyses related to safety, sensation and blinding

The TIS procedure used in the current study was well-tolerated by the participants and no serious adverse events were reported, which is consistent with the growing body of evidence showing that this technique is safe across diverse populations and brain targets (see Demchenko et al., 2025; Vassiliadis et al., 2024 for reviews). In line with previous work (Beanato et al., 2024; Wessel et al., 2023) using similar electrode shape and size (3 $cm^2$), stimulation current intensity (2 mA per channel) and high-frequency control stimulation ($\Delta f = 0$ Hz), we did not observe any differences in sensation ratings (0-3; none, mild, moderate and strong) between active and control stimulation conditions during the habituation phase (statistical results presented in Table S3 and Figure S1A-B). In the post-training sensation questionnaires, the most reported sensations were fatigue/decreased alertness and tingling in both stimulation conditions and experiments (statistical results presented in Table S4 and Figure S1C-D), which is consistent with the sensory effects reported across several TIS studies (Demchenko et al., 2025). Ratings varied across sensation categories but, as during habituation, did not differ between the active and control stimulation conditions in either experiment.

Blinding effectiveness was evaluated at the end of each stimulation session using a debriefing questionnaire adapted from Violante et al. (2023) and Greinacher et al. (2019). Among participants who reported perceiving stimulation, weighted condition-identification scores did not differ from zero in either experiment or condition (Table S5; Figure S2A). Similarly, the participants' beliefs about whether stimulation improved, disrupted, or had no effect on performance did not differ between stimulation conditions and experiments (Table S6; Figure S2B). These results are consistent with previous TIS studies reporting chance-level identification between active and high-frequency control conditions (Beanato et al., 2024; Wessel et al., 2023). Altogether, these results indicate that TIS was well-tolerated and that participants remained blind to the stimulation condition in both experiments.

**Table S3. Stimulation sensation ratings during habituation within each experiment (rating ~ stimulation condition × current intensity) and between experiments (rating ~ experiment × stimulation condition × current intensity).**

| Effect | LR $\chi^2$, df, p |
|---|---|
| **task-TIS (N = 32)** | |
| Stimulation condition | $\chi^2(1) = 0.00, p = .996$ |
| Current intensity | **$\chi^2(3) = 142.78, p < .001$** |
| Stimulation condition × Intensity | $\chi^2(3) = 3.58, p = .311$ |
| **rest-TIS (N = 32)** | |
| Stimulation condition | $\chi^2(1) = 0.14, p = .711$ |
| Current intensity | **$\chi^2(3) = 118.06, p < .001$** |
| Stimulation condition × Intensity | $\chi^2(3) = 6.76, p = .080$ |

| | |
|---|---|
| **Between experiments** | |
| Experiment | $\chi^2(1) = 1.78$, p = .183 |
| Experiment × Stimulation condition | $\chi^2(1) = 0.05$, p = .828 |
| Experiment × Intensity | $\chi^2(3) = 1.66$, p = .646 |
| Experiment × Stimulation condition × Intensity | **$\chi^2(3) = 9.02$, p = .029** |

*Note.* Cumulative link mixed models on ordinal sensation ratings (0–3, absent to strong), with random intercepts for participant and for participant × stimulation condition (visit); fixed effects tested by Type II likelihood-ratio $\chi^2$. Sensation rating increased with current intensity in both experiments, whereas active and control stimulation produced comparable sensations at every current level. Between experiments, only the three-way experiment × stimulation condition × current-intensity interaction was significant and driven by a non-significant difference in intensity ratings between conditions in the rest-TIS experiment (stimulation condition × current intensity within rest-TIS: $\chi^2(3) = 6.76$, p = .080 [task-TIS: $\chi^2(3) = 3.58$, p = .311]; active vs. control contrasts at 0.5, 1.0, 1.5 and 2.0 mA in rest-TIS: b = 1.00, 1.46, 0.13 and −0.71 logits, z = 0.98, 1.86, 0.20 and −1.18, p = .327, .063, .839 and .239, respectively [task-TIS: all p ≥ .185]). Importantly, these marginal differences did not affect blinding (see below). Bold p values indicate p < .05.

**Table S4. Stimulation sensation ratings at post-training within each experiment (rating ~ stimulation condition × sensation category × training phase) and between experiments (rating ~ experiment × stimulation condition × sensation category × training phase).**

| **Effect** | **LR $\chi^2$, df, p** |
|---|---|
| **task-TIS (N = 32)** | |
| Stimulation condition | $\chi^2(1) = 1.61$, p = .205 |
| Sensation category | **$\chi^2(7) = 255.50$, p < .001** |
| Training phase | $\chi^2(1) = 3.85$, p = .050 |
| Stimulation condition × Category | $\chi^2(7) = 4.78$, p = .686 |
| Stimulation condition × Phase | $\chi^2(1) = 0.16$, p = .686 |
| Sensation category × Phase | $\chi^2(7) = 1.50$, p = .982 |
| Stimulation condition × Category × Phase | $\chi^2(7) = 5.40$, p = .611 |
| **rest-TIS (N = 32)** | |
| Stimulation condition | $\chi^2(1) = 0.29$, p = .591 |
| Sensation category | **$\chi^2(7) = 338.52$, p < .001** |
| Training phase | $\chi^2(1) = 2.13$, p = .145 |
| Stimulation condition × Category | $\chi^2(7) = 8.98$, p = .254 |
| Stimulation condition × Phase | $\chi^2(1) = 1.56$, p = .212 |
| Sensation category × Phase | $\chi^2(7) = 5.87$, p = .554 |
| Stimulation condition × Category × Phase | $\chi^2(7) = 2.55$, p = .923 |
| **Between experiments** | |
| Experiment | $\chi^2(1) = 2.44$, p = .118 |
| Experiment × Stimulation condition | $\chi^2(1) = 1.45$, p = .228 |
| Experiment × Category | $\chi^2(7) = 12.26$, p = .092 |
| Experiment × Phase | $\chi^2(1) = 0.13$, p = .721 |
| Experiment × Stimulation condition × Category × Phase | $\chi^2(7) = 5.92$, p = .550 |

*Note.* Cumulative link mixed models on ordinal ratings (0–3), with random intercepts for participant and for participant × stimulation condition (visit); fixed effects tested by Type II likelihood-ratio $\chi^2$ (nested models). Ratings differed between sensation categories in both experiments, with fatigue/decreased alertness and tingling the most reported (Figure S1C–D), but did not differ between the active and control conditions, and stimulation condition did not interact with sensation category, training phase or experiment. Bold p values indicate p < .05.

**Table S5. Condition identification (weighted score) within each experiment (Wilcoxon signed-rank tests against zero and stimulation-condition cumulative link mixed models) and between experiments (experiment × stimulation condition cumulative link mixed model).**

| **Condition / Effect** | **Mdn [IQR]; M ± SD; test statistic, p** |
|---|---|
| **task-TIS (N = 32)** | |
| Active (n = 32) | Mdn = 3 [-5.25, 6.125]; M ± SD = 0.80 ± 6.46; V = 300, p = .506 |
| Control (n = 32) | Mdn = 4.25 [-6, 6.375]; M ± SD = 0.88 ± 6.89; V = 305.5, p = .442 |
| Active vs Control (stimulation condition) | $\chi^2(1) = 0.03$, p = .872 |
| **rest-TIS (N = 32)** | |
| Active (n = 32) | Mdn = -0.5 [-6.5, 4.25]; M ± SD = -1.02 ± 6.38; V = 190, p = .259 |
| Control (n = 31) | Mdn = -2 [-7, 7]; M ± SD = -0.27 ± 6.70; V = 240, p = .883 |
| Active vs Control (stimulation condition) | $\chi^2(1) = 0.57$, p = .448 |
| **Between experiments: 2 (Experiment) × 2 (Stimulation condition)** | |
| Experiment | $\chi^2(1) = 1.82$, p = .177 |
| Experiment × Stimulation condition | $\chi^2(1) = 0.13$, p = .720 |

*Note*. The weighted condition-identification score is the guessed condition (active = +1, control = −1) multiplied by the confidence rating (1 to 10; not confident to extremely confident); a mean of zero corresponds to chance-level identification. Cumulative link mixed models (CLMMs; logit link) were conducted including a random intercept for participant; fixed effects were tested with likelihood-ratio $\chi^2$ tests (within each experiment: stimulation condition; between experiments: experiment, stimulation condition and their interaction). No participant-by-visit random intercept was included because each participant provided a single score per visit, so this term would be confounded with the residual. Deviation from zero within each cell was tested with Wilcoxon signed-rank tests (V statistic). Medians (Mdn) with interquartile ranges [Q1, Q3] and means ± SD are given for description. One session (E54, rest-TIS control) had no debriefing data and is not included.

**Table S6. Perceived effect of stimulation on performance within each experiment (Fisher's exact test) and between experiments (Cochran–Mantel–Haenszel test).**

| **Condition / Effect** | **Improved / Disrupted / No effect** |
|---|---|
| **task-TIS (N = 32)** | |
| Active | 8 / 5 / 16 |
| Control | 9 / 7 / 11 |
| Active vs Control | Fisher's exact p = .573 |
| **rest-TIS (N = 32)** | |
| Active | 8 / 5 / 14 |
| Control | 8 / 6 / 14 |
| Active vs Control | Fisher's exact p = 1.000 |
| **Between experiments** | |
| Experiment | $\chi^2(1) = 0.01$, p = .905 |
| Experiment × Stimulation condition | $\chi^2(1) = 0.02$, p = .892 |

*Note.* Frequencies of participants reporting that stimulation improved, disrupted, or had no effect on performance. Within each experiment the distribution was compared between conditions with a Fisher's exact test, which does not yield a test statistic; the two experiments were compared with a Cochran–Mantel–Haenszel test.

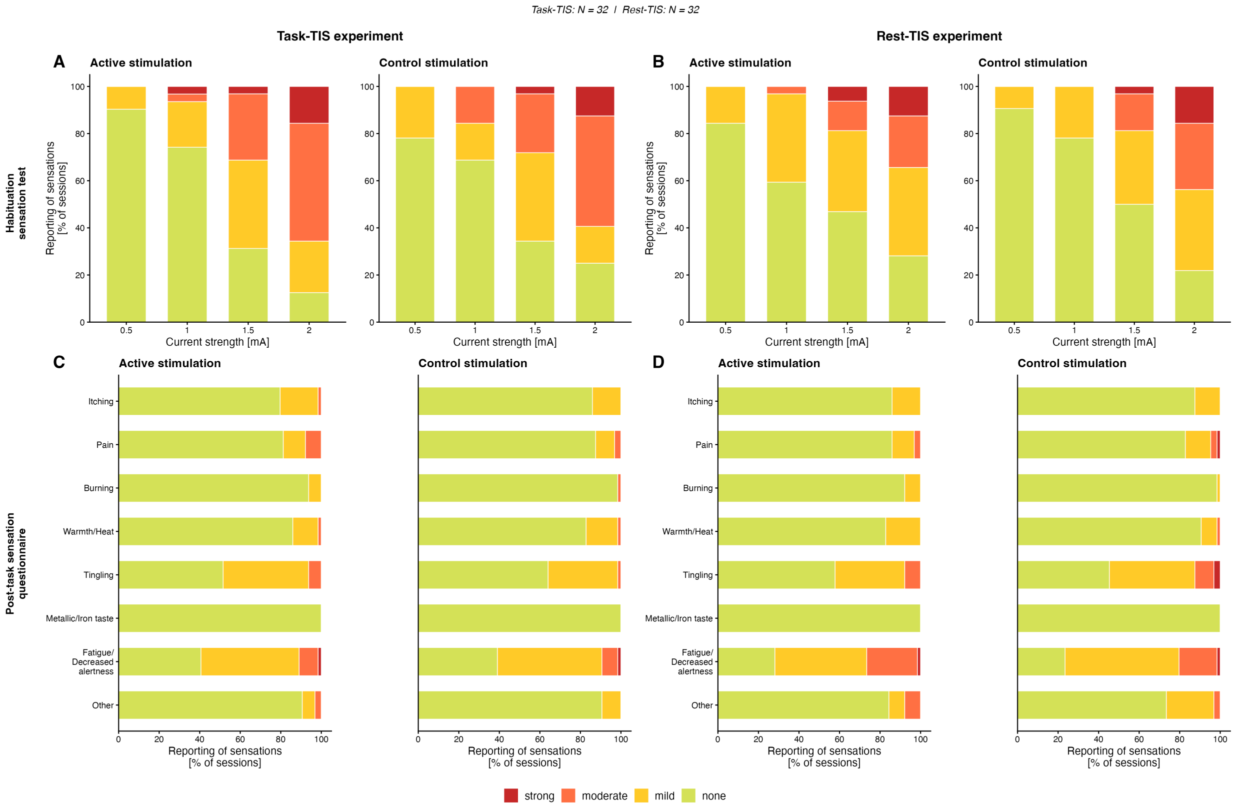

**Figure S1. Stimulation-related sensations during habituation (A-B) and after each training session (C-D) in the task-TIS and rest-TIS experiments for the active and control stimulation conditions.** Frequency of reported sensations separated by their strength (none, mild, moderate, and strong) are depicted for the tested current intensity levels (0.5, 1.0, 1.5, 2 mA per stimulation channel) during the habituation sensation (A-B) and for the different sensation category (itching, pain, burning, warmth, tingling, metallic taste, fatigue and other) after training (pooled across the two training phases of each visit) (C–D).

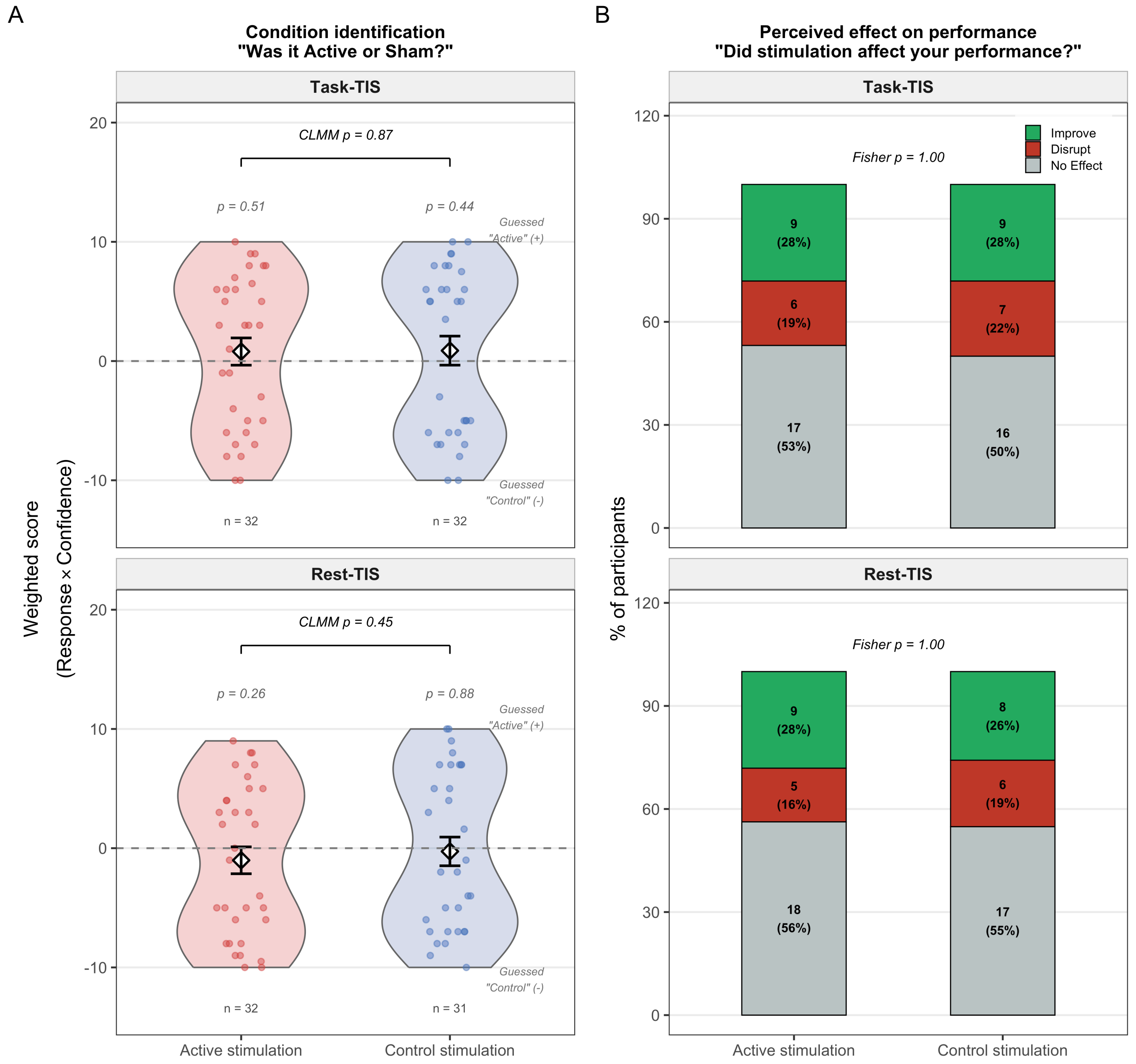

**Figure S2. Stimulation condition identification (A) and perceived effect of stimulation on performance (B) in the task-TIS and rest-TIS experiments for the two different stimulation conditions (active vs control TIS). A.** A weighted score related to the question "Do you think you received active or control stimulation?" was computed by multiplying the response type (yes = +1; no = −1) by the confidence rating (1 to 10) such that higher positive scores indicate greater confidence in having received active stimulation while higher negative scores indicate greater confidence in having received control stimulation. **B.** Percentage of participants indicating they thought that stimulation improved / disrupt / had no effect on their performance.

### 3. Exploratory electric field modeling

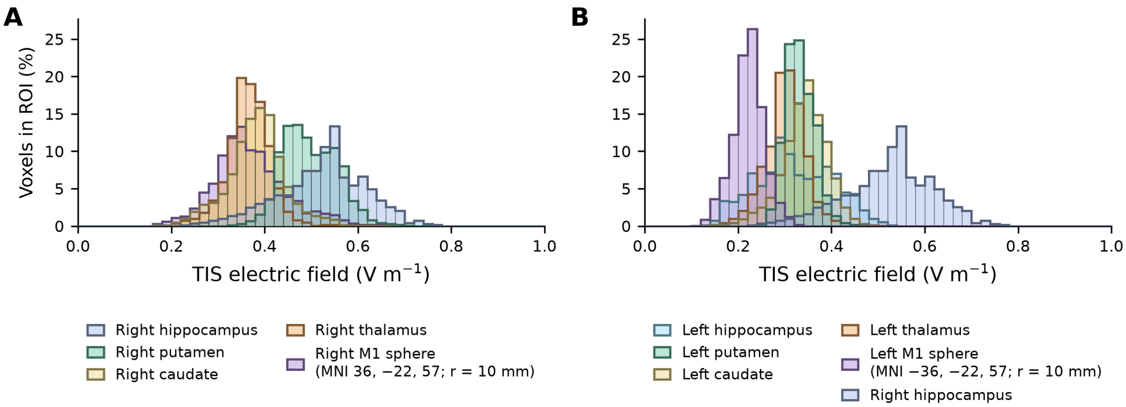

**Figure S3. TIS electric field exposure in the right hippocampus and in other task-relevant regions (putamen, caudate, thalamus and the M1 hand area).** The TIS-induced electric fields were simulated on the MIDA head model (Iacono et al., 2015) in Sim4Life.lite using the experimental montage FT8–Fp1 / TP8–P7 (2 mA per channel), with anisotropic (DTI-based) conductivity in white matter and midbrain (see Methods). Histograms show the field distribution, expressed as the percentage of voxels within each region of interest (ROI), in the right hippocampus and in other task-relevant brain regions in the right (**A**) and left (**B**) hemispheres (note that the right hippocampus is also presented in panel B for reference). The deep structures were defined from the corresponding MIDA tissue labels and the M1 hand area as the cortical grey-matter voxels within a 10-mm-radius sphere centered on MNI coordinates (±36, −22, 57 mm) (see Methods). Median field per ROI, in V $m^{-1}$: right hippocampus 0.53, right putamen 0.49, right caudate 0.38, right thalamus 0.37, right M1 0.36; left hippocampus 0.31, left putamen 0.33, left caudate 0.35, left thalamus 0.30, left M1 0.22.

### 4. Supplemental results on performance speed

**Table S7. Performance speed (cIRI): full repeated-measures ANOVA models for training and the post-training test, within and between experiments.**

| Effect | F, df, p, $\eta^2p$ |
|---|---|
| **A. Training — 2 (Training phase) × 20 (Block) × 2 (Stimulation condition)** | |
| **task-TIS (N = 32)** | |
| Training phase | **F(1.00, 31.00) = 163.94, p < .001, $\eta^2p$ = 0.841** |
| Block | **F(3.59, 111.35) = 77.50, p < .001, $\eta^2p$ = 0.714** |
| Stimulation condition | F(1.00, 31.00) = 0.15, p = .697, $\eta^2p$ = 0.005 |
| Training phase × Stimulation condition | F(1.00, 31.00) = 0.26, p = .616, $\eta^2p$ = 0.008 |
| Block × Stimulation condition | F(3.96, 122.84) = 1.06, p = .381, $\eta^2p$ = 0.033 |
| Training phase × Block × Stimulation condition | F(4.91, 152.33) = 0.88, p = .497, $\eta^2p$ = 0.027 |
| **rest-TIS (N = 32)** | |
| Training phase | **F(1.00, 31.00) = 170.82, p < .001, $\eta^2p$ = 0.846** |
| Block | **F(3.49, 108.25) = 80.86, p < .001, $\eta^2p$ = 0.723** |
| Stimulation condition | F(1.00, 31.00) = 0.04, p = .836, $\eta^2p$ = 0.001 |
| Training phase × Stimulation condition | F(1.00, 31.00) = 0.85, p = .365, $\eta^2p$ = 0.027 |
| Block × Stimulation condition | F(3.99, 123.58) = 0.33, p = .856, $\eta^2p$ = 0.011 |
| Training phase × Block × Stimulation condition | F(3.73, 115.52) = 0.44, p = .768, $\eta^2p$ = 0.014 |
| **Between experiments (N = 64)** | |

| | |
|---|---|
| Experiment | F(1.00, 62.00) = 0.67, p = .416, η²p = 0.011 |
| Experiment × Training phase | F(1.00, 62.00) = 0.13, p = .724, η²p = 0.002 |
| Experiment × Block | F(3.76, 233.27) = 0.65, p = .620, η²p = 0.010 |
| Experiment × Stimulation condition | F(1.00, 62.00) = 0.03, p = .858, η²p = 0.001 |
| Experiment × Training phase × Block × Stimulation condition | F(4.77, 295.43) = 0.44, p = .814, η²p = 0.007 |
| **B. Post-training test — 4 (Block) × 2 (Stimulation condition)** | |
| **task-TIS (N = 32)** | |
| Stimulation condition | F(1.00, 31.00) = 0.01, p = .919, η²p = 0.000 |
| Block | F(2.50, 77.64) = 0.44, p = .693, η²p = 0.014 |
| Block × Stimulation condition | F(1.88, 58.35) = 0.11, p = .883, η²p = 0.004 |
| **rest-TIS (N = 32)** | |
| Stimulation condition | F(1.00, 31.00) = 3.00, p = .093, η²p = 0.088 |
| Block | F(2.57, 79.70) = 1.66, p = .189, η²p = 0.051 |
| Block × Stimulation condition | F(2.41, 74.60) = 0.15, p = .894, η²p = 0.005 |
| **Between experiments (N = 64)** | |
| Experiment | F(1.00, 62.00) = 0.89, p = .350, η²p = 0.014 |
| Experiment × Block | F(2.55, 158.02) = 0.02, p = .991, η²p = 0.000 |
| Experiment × Stimulation condition | F(1.00, 62.00) = 1.38, p = .245, η²p = 0.022 |
| Experiment × Block × Stimulation condition | F(2.38, 147.41) = 0.04, p = .976, η²p = 0.001 |

*Note.* Greenhouse–Geisser correction applied to within-subject factors with more than one numerator df. Performance improved across blocks and training phases in both experiments; there was no main effect or interaction involving stimulation condition, and no between-experiment difference in the condition effect. There was no effect of stimulation condition on post-training data in either experiment.

### 5. Supplemental results on performance accuracy

**Table S8. Performance accuracy (% correct transitions): full repeated-measures ANOVA models for training and the post-training test, and macro-offline changes, within and between experiments.**

| Effect | F, df, p, η²p |
|---|---|
| **A. Training — 2 (Training phase) × 20 (Block) × 2 (Stimulation condition)** | |
| **task-TIS (N = 32)** | |
| Training phase | **F(1.00, 31.00) = 20.53, p < .001, η²p = 0.398** |
| Block | F(10.60, 328.65) = 0.96, p = .481, η²p = 0.030 |
| Stimulation condition | F(1.00, 31.00) = 0.59, p = .450, η²p = 0.019 |
| Training phase × Block | F(10.23, 317.20) = 1.71, p = .075, η²p = 0.052 |
| Training phase × Stimulation condition | F(1.00, 31.00) = 0.54, p = .467, η²p = 0.017 |
| Block × Stimulation condition | F(10.59, 328.40) = 1.05, p = .398, η²p = 0.033 |
| Training phase × Block × Stimulation condition | F(10.74, 332.97) = 0.84, p = .596, η²p = 0.026 |
| **rest-TIS (N = 32)** | |
| Training phase | **F(1.00, 31.00) = 13.96, p < .001, η²p = 0.310** |
| Block | F(7.66, 237.59) = 1.65, p = .115, η²p = 0.051 |
| Stimulation condition | F(1.00, 31.00) = 2.68, p = .112, η²p = 0.080 |
| Training phase × Block | F(8.73, 270.59) = 1.13, p = .342, η²p = 0.035 |
| Training phase × Stimulation condition | F(1.00, 31.00) = 1.52, p = .227, η²p = 0.047 |
| Block × Stimulation condition | F(8.54, 264.75) = 1.22, p = .283, η²p = 0.038 |
| Training phase × Block × Stimulation condition | F(8.29, 256.88) = 0.94, p = .487, η²p = 0.029 |
| **Between experiments (N = 64)** | |

| | |
|---|---|
| Experiment | $F(1.00, 62.00) = 0.09$, $p = .769$, $\eta^2p = 0.001$ |
| Experiment × Training phase | $F(1.00, 62.00) = 0.88$, $p = .353$, $\eta^2p = 0.014$ |
| Experiment × Block | $F(11.15, 691.56) = 0.70$, $p = .739$, $\eta^2p = 0.011$ |
| Experiment × Stimulation condition | $F(1.00, 62.00) = 3.14$, $p = .081$, $\eta^2p = 0.048$ |
| Experiment × Training phase × Block × Stimulation condition | $F(12.98, 804.60) = 0.97$, $p = .477$, $\eta^2p = 0.015$ |
| **B. Post-training test — 4 (Block) × 2 (Stimulation condition)** | |
| **task-TIS (N = 32)** | |
| Stimulation condition | $F(1, 31) = 0.00$, $p = .982$, $\eta^2p = 0.000$ |
| Block | $F(3, 93) = 2.67$, $p = .052$, $\eta^2p = 0.079$ |
| Block × Stimulation condition | $F(3, 93) = 0.61$, $p = .610$, $\eta^2p = 0.019$ |
| **rest-TIS (N = 32)** | |
| Stimulation condition | $F(1, 31) = 3.15$, $p = .086$, $\eta^2p = 0.092$ |
| Block | $F(3, 93) = 0.90$, $p = .447$, $\eta^2p = 0.028$ |
| Block × Stimulation condition | $F(3, 93) = 2.39$, $p = .074$, $\eta^2p = 0.072$ |
| **Between experiments (N = 64)** | |
| Experiment | $F(1, 62) = 0.00$, $p = .991$, $\eta^2p = 0.000$ |
| Experiment × Block | **$F(3, 186) = 3.30$, $p = .022$, $\eta^2p = 0.051$** |
| Experiment × Stimulation condition | $F(1, 62) = 2.06$, $p = .156$, $\eta^2p = 0.032$ |
| Experiment × Block × Stimulation condition | $F(3, 186) = 2.34$, $p = .075$, $\eta^2p = 0.036$ |
| **C. Macro-offline changes in performance** | |
| **task-TIS (N = 32)** | |
| Active vs Control | $t(31) = 0.57$, $p = .570$, $d = 0.10$ |
| **rest-TIS (N = 32)** | |
| Active vs Control | $t(31) = -1.53$, $p = .136$, $d = -0.27$ |
| **Between experiments (N = 64)** | |
| Experiment | $F(1, 62) = 0.30$, $p = .587$, $\eta^2p = 0.005$ |
| Experiment × Stimulation condition | $F(1, 62) = 1.92$, $p = .171$, $\eta^2p = 0.030$ |

*Note.* Greenhouse–Geisser correction applied to within-subject factors with more than one numerator df. Accuracy was high and increased from Training 1 to Training 2 phases in both conditions and experiments. There was no main effect or interaction involving stimulation condition on training or post-training test accuracy in either experiment, and no between-experiment difference including the stimulation condition effect. Macro-offline changes in performance accuracy did not differ between stimulation conditions in either experiment, and there was no between-experiment difference.

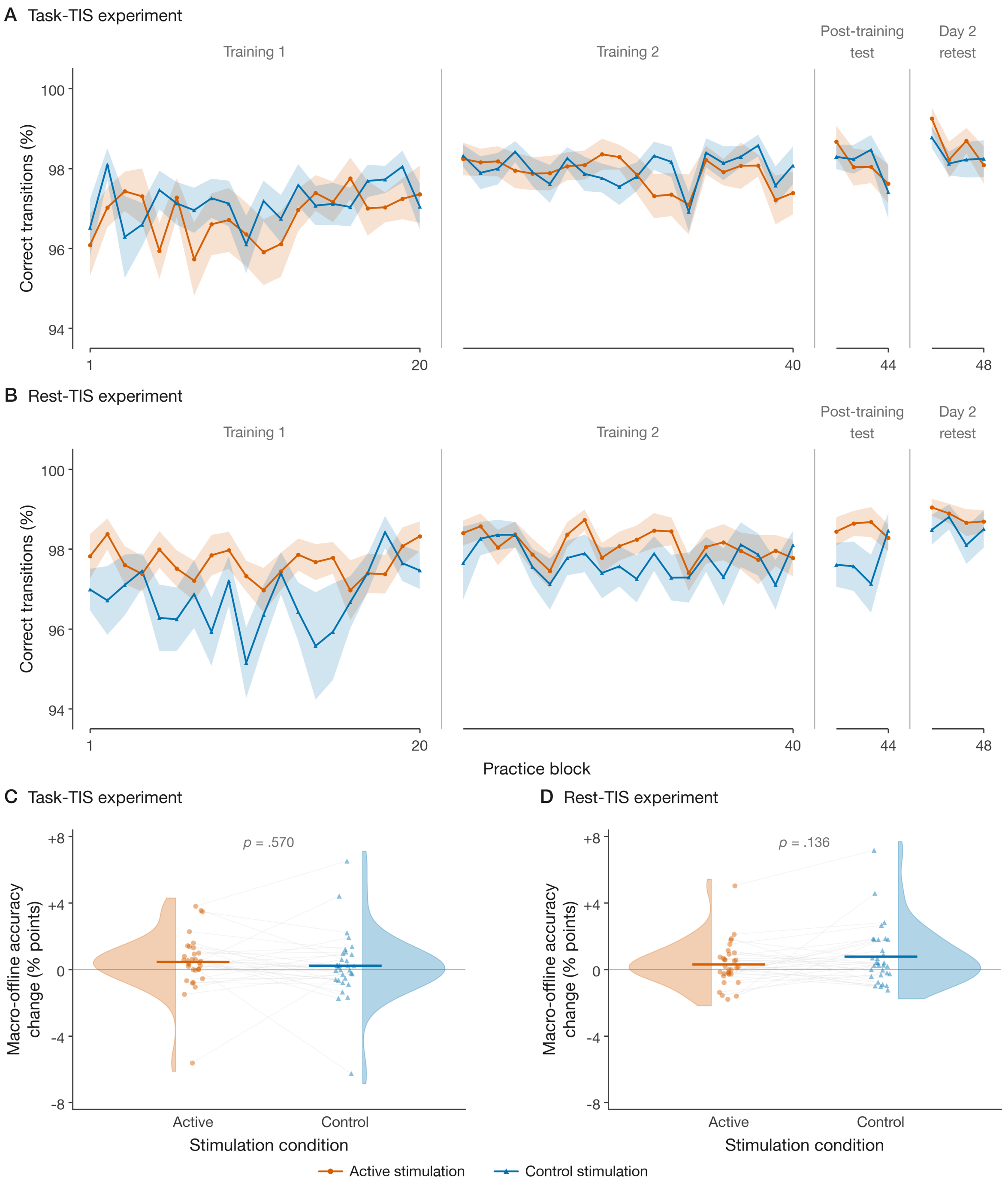

**Figure S4. Performance accuracy.** ***A-B***. Percentage of correct transitions across the 40 training blocks, the post-training test and the Day 2 retest, for the active (orange) and control (blue) stimulation conditions in the task-TIS (A) and rest-TIS (B) experiments. Lines and shaded areas represent means ± SEM across participants. **C-D.** Macro-offline changes in performance accuracy for the active and control stimulation conditions in the task-TIS (C) and rest-TIS (D) experiments. Dots represent individual participants, horizontal bars the condition means, and half-violins the kernel density estimate; p values are from paired t-tests (active vs. control). Corresponding statistics are reported in Supplemental Table S8.

### 6. Spatial and motor representations as well as transfer effects

In both the task- and rest-TIS experiments, hippocampal stimulation failed to modulate performance on the spatial (or motoric) representations of the learned sequence (Supplemental Tables S9 and S10; Supplemental Figures S5 and S6). This was unexpected as previous work from our group has established that the hippocampus is critically involved in encoding the allocentric, spatial representation of learned motor sequences during initial learning (Albouy et al., 2013, 2015; Dolfen et al., 2024), and that these spatial representations are spontaneously replayed in the hippocampus during post-learning rest (King et al., 2022). One potential explanation for this null effect could be related to the timing of the representation testing in the current study. Specifically, spatial and motor representations of the task were tested on Day 2 (as opposed to immediately after learning as in our prior research (Albouy et al., 2013, 2015) and it is possible that the longer consolidation interval (including a full night of sleep) may have attenuated the potential effect of stimulation on performance (see main discussion for reflections on potential transient effects).

**Table S9. Day 2 representation and transfer tests: performance speed.**

| Effect | F, df, p, $\eta^2p$ |
|---|---|
| **A. Representation tests — 2 (Representation) × 4 (Block) × 2 (Stimulation condition)** | |
| **task-TIS (N = 32)** | |
| Representation | $F(1, 31) = 0.42$, $p = .523$, $\eta^2p = 0.013$ |
| Block | **$F(1.25, 38.80) = 53.94$, $p < .001$, $\eta^2p = 0.635$** |
| Stimulation condition | $F(1, 31) = 0.02$, $p = .882$, $\eta^2p = 0.001$ |
| Representation × Block | $F(1.93, 59.89) = 0.36$, $p = .694$, $\eta^2p = 0.011$ |
| Representation × Stimulation condition | $F(1, 31) = 0.20$, $p = .658$, $\eta^2p = 0.006$ |
| Block × Stimulation condition | $F(1.81, 56.07) = 1.05$, $p = .349$, $\eta^2p = 0.033$ |
| Representation × Block × Stimulation condition | $F(1.86, 57.65) = 0.78$, $p = .455$, $\eta^2p = 0.025$ |
| **rest-TIS (N = 32)** | |
| Representation | $F(1, 31) = 4.09$, $p = .052$, $\eta^2p = 0.116$ |
| Block | **$F(1.25, 38.59) = 89.96$, $p < .001$, $\eta^2p = 0.744$** |
| Stimulation condition | $F(1, 31) = 0.00$, $p = .996$, $\eta^2p = 0.000$ |
| Representation × Block | $F(1.81, 56.23) = 0.27$, $p = .740$, $\eta^2p = 0.009$ |
| Representation × Stimulation condition | $F(1, 31) = 0.18$, $p = .671$, $\eta^2p = 0.006$ |
| Block × Stimulation condition | $F(1.62, 50.32) = 0.14$, $p = .829$, $\eta^2p = 0.004$ |
| Representation × Block × Stimulation condition | $F(1.74, 53.89) = 1.29$, $p = .281$, $\eta^2p = 0.040$ |
| **Between experiments (N = 64)** | |
| Experiment | $F(1, 62) = 1.74$, $p = .193$, $\eta^2p = 0.027$ |
| Experiment × Representation | $F(1, 62) = 1.04$, $p = .312$, $\eta^2p = 0.016$ |
| Experiment × Block | $F(1.30, 80.83) = 0.14$, $p = .773$, $\eta^2p = 0.002$ |
| Experiment × Stimulation condition | $F(1, 62) = 0.01$, $p = .907$, $\eta^2p = 0.000$ |
| Experiment × Representation × Block × Stimulation condition | $F(1.84, 114.32) = 1.76$, $p = .179$, $\eta^2p = 0.028$ |
| **B. Transfer indices — 2 (Transfer type) × 2 (Stimulation condition)** | |
| **task-TIS (N = 32)** | |
| Transfer type | $F(1, 31) = 0.42$, $p = .523$, $\eta^2p = 0.013$ |
| Stimulation condition | $F(1, 31) = 0.00$, $p = .959$, $\eta^2p = 0.000$ |
| Transfer type × Stimulation condition | $F(1, 31) = 0.20$, $p = .658$, $\eta^2p = 0.006$ |
| **rest-TIS (N = 32)** | |
| Transfer type | $F(1, 31) = 4.09$, $p = .052$, $\eta^2p = 0.116$ |
| Stimulation condition | $F(1, 31) = 0.01$, $p = .927$, $\eta^2p = 0.000$ |
| Transfer type × Stimulation condition | $F(1, 31) = 0.18$, $p = .671$, $\eta^2p = 0.006$ |

| | |
|---|---|
| **Between experiments (N = 64)** | |
| Experiment | F(1, 62) = 0.63, p = .429, η²p = 0.010 |
| Experiment × Transfer type | F(1, 62) = 1.04, p = .312, η²p = 0.016 |
| Experiment × Stimulation condition | F(1, 62) = 0.01, p = .922, η²p = 0.000 |
| Experiment × Transfer type × Stimulation condition | F(1, 62) = 0.38, p = .538, η²p = 0.006 |

*Note.* There was no effect of stimulation condition on performance speed for the spatial or motor representations of the task, in either experiment or between experiments. Transfer indices (novel sequence minus spatial/motor representation, see methods) based on the performance speed measure did not differ between the active and control stimulation conditions in either experiment or between experiments.

**Table S10. Day 2 representation and transfer tests: performance accuracy.**

| Effect | F, df, p, η²p |
|---|---|
| **A. Representation tests — 2 (Representation) × 4 (Block) × 2 (Stimulation condition)** | |
| **task-TIS (N = 32)** | |
| Representation | F(1, 31) = 0.00, p = .990, η²p = 0.000 |
| Block | F(1.60, 49.71) = 0.44, p = .602, η²p = 0.014 |
| Stimulation condition | F(1, 31) = 2.01, p = .166, η²p = 0.061 |
| Representation × Block | F(1.73, 53.65) = 1.86, p = .170, η²p = 0.057 |
| Representation × Stimulation condition | F(1, 31) = 1.19, p = .283, η²p = 0.037 |
| Block × Stimulation condition | F(1.83, 56.61) = 2.67, p = .082, η²p = 0.079 |
| Representation × Block × Stimulation condition | F(1.84, 56.91) = 0.20, p = .800, η²p = 0.006 |
| **rest-TIS (N = 32)** | |
| Representation | F(1, 31) = 0.00, p = .994, η²p = 0.000 |
| Block | F(2.76, 85.41) = 1.11, p = .347, η²p = 0.035 |
| Stimulation condition | F(1, 31) = 3.84, p = .059, η²p = 0.110 |
| Representation × Block | F(2.17, 67.38) = 1.82, p = .168, η²p = 0.055 |
| Representation × Stimulation condition | F(1, 31) = 0.64, p = .429, η²p = 0.020 |
| Block × Stimulation condition | F(2.82, 87.38) = 0.73, p = .531, η²p = 0.023 |
| Representation × Block × Stimulation condition | F(2.90, 89.82) = 0.83, p = .479, η²p = 0.026 |
| **Between experiments (N = 64)** | |
| Experiment | F(1, 62) = 0.28, p = .600, η²p = 0.004 |
| Experiment × Representation | F(1, 62) = 0.00, p = .994, η²p = 0.000 |
| Experiment × Block | F(1.98, 122.95) = 0.61, p = .545, η²p = 0.010 |
| Experiment × Stimulation condition | F(1, 62) = 0.01, p = .942, η²p = 0.000 |
| Experiment × Representation × Block × Stimulation condition | F(2.21, 137.01) = 0.61, p = .561, η²p = 0.010 |
| **B. Transfer indices — 2 (Transfer type) × 2 (Stimulation condition)** | |
| **task-TIS (N = 32)** | |
| Transfer type | F(1, 31) = 0.00, p = .990, η²p = 0.000 |
| Stimulation condition | **F(1, 31) = 8.61, p = .006, η²p = 0.217** |
| Transfer type × Stimulation condition | F(1, 31) = 1.19, p = .283, η²p = 0.037 |
| **rest-TIS (N = 32)** | |
| Transfer type | F(1, 31) = 0.00, p = .994, η²p = 0.000 |
| Stimulation condition | F(1, 31) = 0.00, p = .979, η²p = 0.000 |
| Transfer type × Stimulation condition | F(1, 31) = 0.64, p = .429, η²p = 0.020 |
| **Between experiments (N = 64)** | |
| Experiment | F(1, 62) = 0.26, p = .610, η²p = 0.004 |
| Experiment × Transfer type | F(1, 62) = 0.00, p = .994, η²p = 0.000 |

| | |
|---|---|
| Experiment × Stimulation condition | **F(1, 62) = 4.87, p = .031, $\eta^2p$ = 0.073** |
| Experiment × Transfer type × Stimulation condition | F(1, 62) = 1.74, p = .192, $\eta^2p$ = 0.027 |

*Note.* Greenhouse–Geisser correction applied to within-subject factors with more than one numerator df. **A.** During the representation test runs, accuracy was higher in the active than in the control stimulation condition when collapsing across experiments and representations, with no condition effect within either experiment and no other significant effect involving stimulation condition. The experiment × representation × block interaction was driven by an experiment × block effect in the motor representation (F(2.21, 137.25) = 3.56, p = .027, $\eta^2p$ = 0.054). Follow-up tests showed no significant effect of block within either experiment (task-TIS: p = .178; rest-TIS: p = .124) and no difference between experiments at any block (all p ≥ .078), indicating differences in block-to-block fluctuations in accuracy on the motor representation between experiments. **B.** With respect to the transfer indices, the stimulation condition effect was significant in the task-TIS experiment whereby transfer was higher in the control than in the active stimulation condition. This effect was greater in the task-TIS compared to the rest-TIS experiment.

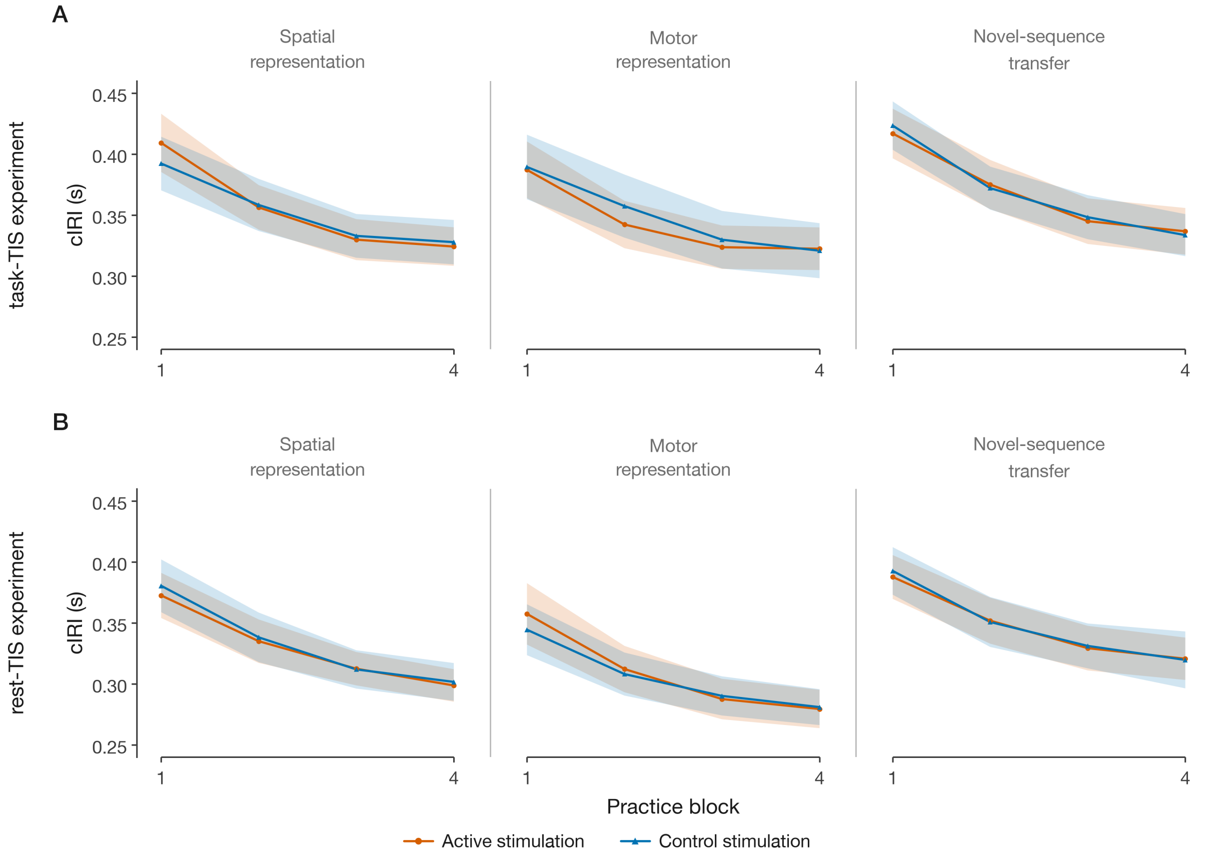

**Figure S5. Performance speed on spatial and motor representations as well on the novel sequence performed on Day 2** for the task-TIS (A) and rest-TIS (B) experiments (n = 32 in each experiment) and for the active (orange) and control (blue) stimulation conditions. Performance speed, measured as the inter-response interval for correct key presses (cIRI, in s), is plotted for each block of practice of the spatial representation, motor representation and novel sequence (4 blocks in each condition). Lower values indicate faster performance. Shaded regions represent SEM. There was no effect of stimulation condition on these tasks in either experiment.

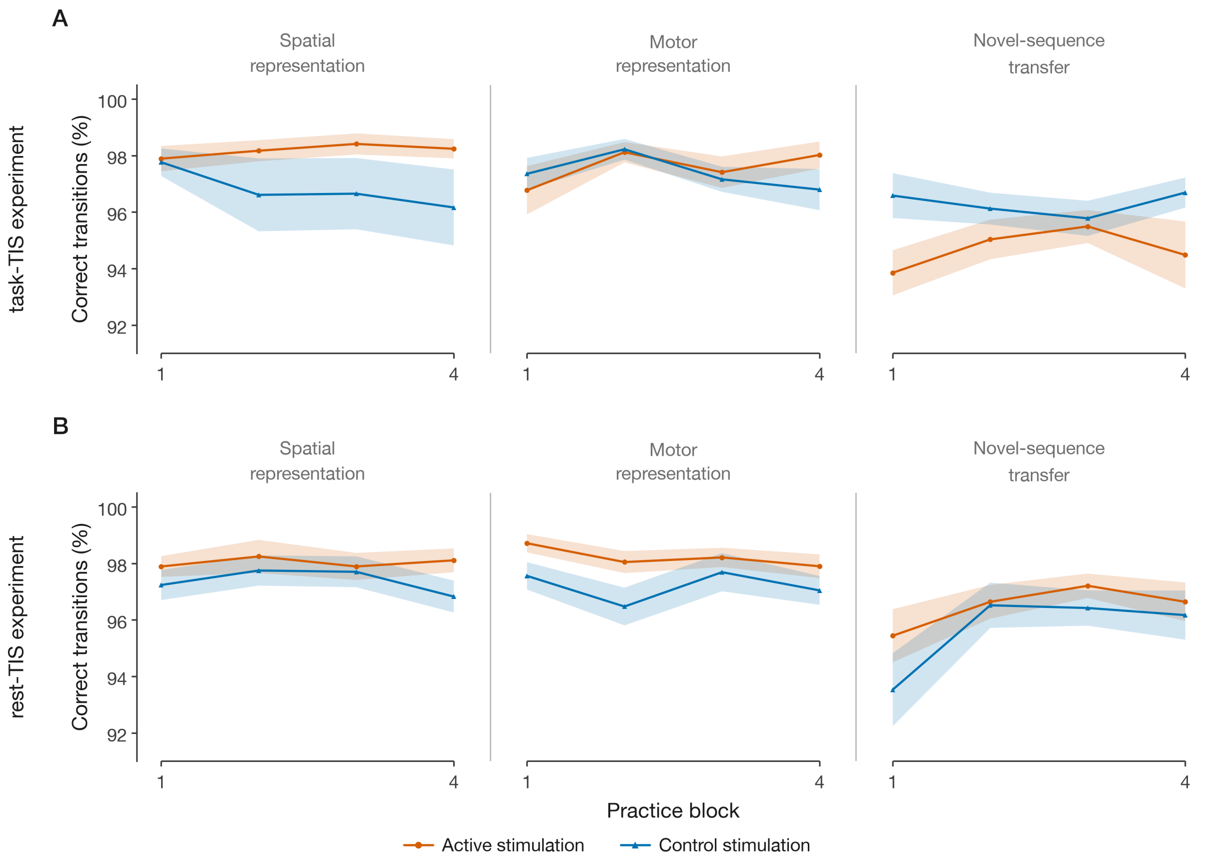

**Figure S6. Performance accuracy on spatial and motor representations as well on the novel sequence performed on Day 2** for the task-TIS (A) and rest-TIS (B) experiments (n = 32 in each experiment) and for the active (orange) and control (blue) stimulation conditions. Performance accuracy, measured as the percentage of correct transitions, is plotted for each block of practice of the spatial representation, motor representation and novel sequence (4 blocks in each condition). Higher values indicate more accurate performance. Shaded regions represent SEM. There was no effect of stimulation condition on accuracy for the spatial and motor representation tests in either experiment.

### 7. Supplemental micro timescale results

**Table S11. Micro-timescale changes in performance speed: full models, simple effects, with and without the participant showing extreme micro-offline gains.**

| Effect | F / t, df, p, $\eta^2p$ / d |
|---|---|
| **A. 2 (Micro type) × 2 (Stimulation condition), with follow-up paired comparisons** | |
| **task-TIS (N = 32)** | |
| Micro type | **$F(1.00, 31.00) = 92.50$, $p < .001$, $\eta^2p = 0.749$** |
| Stimulation condition | $F(1.00, 31.00) = 1.47$, $p = .234$, $\eta^2p = 0.045$ |
| Micro type × Stimulation condition | $F(1.00, 31.00) = 0.18$, $p = .677$, $\eta^2p = 0.006$ |
| **rest-TIS (N = 32)** | |
| Micro type | **$F(1.00, 31.00) = 31.68$, $p < .001$, $\eta^2p = 0.505$** |
| Stimulation condition | $F(1.00, 31.00) = 0.21$, $p = .646$, $\eta^2p = 0.007$ |
| Micro type × Stimulation condition | **$F(1.00, 31.00) = 8.67$, $p = .006$, $\eta^2p = 0.219$** |
| Micro-offline: Active vs Control | **$t(31) = -3.10$, $p = .004$, $d = -0.55$** |

| | |
|---|---|
| Micro-online: Active vs Control | **t(31) = 2.77, p = .009, d = 0.49** |
| **Between experiments (N = 64)** | |
| Experiment | F(1.00, 62.00) = 0.43, p = .515, $\eta^2p$ = 0.007 |
| Experiment × Micro type | F(1.00, 62.00) = 0.32, p = .576, $\eta^2p$ = 0.005 |
| Experiment × Stimulation condition | F(1.00, 62.00) = 1.20, p = .279, $\eta^2p$ = 0.019 |
| Experiment × Micro type × Stimulation condition | **F(1.00, 62.00) = 5.32, p = .024, $\eta^2p$ = 0.079** |
| **B. 2 (Micro type) × 2 (Stimulation condition), with follow-up paired comparisons excluding the extreme participant** | |
| **rest-TIS (N = 31)** | |
| Micro type | **F(1.00, 30.00) = 48.33, p < .001, $\eta^2p$ = 0.617** |
| Stimulation condition | F(1.00, 30.00) = 0.07, p = .791, $\eta^2p$ = 0.002 |
| Micro type × Stimulation condition | **F(1.00, 30.00) = 7.30, p = .011, $\eta^2p$ = 0.196** |
| Micro-offline: Active vs Control | **t(30) = -2.85, p = .008, d = -0.51** |
| Micro-online: Active vs Control | **t(30) = 2.54, p = .017, d = 0.46** |
| **Between experiments (N = 63)** | |
| Experiment | F(1.00, 61.00) = 0.56, p = .456, $\eta^2p$ = 0.009 |
| Experiment × Micro type | F(1.00, 61.00) = 2.13, p = .149, $\eta^2p$ = 0.034 |
| Experiment × Stimulation condition | F(1.00, 61.00) = 0.42, p = .520, $\eta^2p$ = 0.007 |
| Experiment × Micro type × Stimulation condition | **F(1.00, 61.00) = 4.29, p = .043, $\eta^2p$ = 0.066** |
| **C. Exploratory analysis on Training 1 only (blocks 1–20, i.e., 19 inter-practice rest periods): 2 (Micro type) × 2 (Stimulation condition), with follow-up paired comparisons** | |
| **task-TIS (N = 32)** | |
| Micro type | **F(1.00, 31.00) = 100.25, p < .001, $\eta^2p$ = 0.764** |
| Stimulation condition | F(1.00, 31.00) = 1.39, p = .247, $\eta^2p$ = 0.043 |
| Micro type × Stimulation condition | F(1.00, 31.00) = 0.55, p = .464, $\eta^2p$ = 0.017 |
| **rest-TIS (N = 32)** | |
| Micro type | **F(1.00, 31.00) = 33.04, p < .001, $\eta^2p$ = 0.516** |
| Stimulation condition | F(1.00, 31.00) = 0.01, p = .940, $\eta^2p$ = 0.000 |
| Micro type × Stimulation condition | **F(1.00, 31.00) = 5.70, p = .023, $\eta^2p$ = 0.155** |
| Micro-offline: Active vs Control | **t(31) = -2.50, p = .018, d = -0.44** |
| Micro-online: Active vs Control | **t(31) = 2.23, p = .033, d = 0.39** |
| **D. Same as C, excluding the extreme participant** | |
| **rest-TIS (N = 31)** | |
| Micro type | **F(1.00, 30.00) = 42.76, p < .001, $\eta^2p$ = 0.588** |
| Stimulation condition | F(1.00, 30.00) = 0.52, p = .478, $\eta^2p$ = 0.017 |
| Micro type × Stimulation condition | **F(1.00, 30.00) = 4.63, p = .040, $\eta^2p$ = 0.134** |
| Micro-offline: Active vs Control | **t(30) = -2.33, p = .027, d = -0.42** |
| Micro-online: Active vs Control | t(30) = 1.95, p = .061, d = 0.35 |

*Exploratory analyses on early learning.* Based on earlier research suggesting that fast consolidation processes are preferentially observed during early learning stages (Bönstrup et al., 2019), we performed exploratory analyses on the micro gains accumulated over Training 1 phase only. Micro-online and micro-offline changes in performance summed over Training 1 were analyzed with the same within-experiment models as in the main analyses (Table S11C and D). No stimulation effect was observed in the task-TIS experiment, whereas in the rest-TIS experiment, the micro-learning type × stimulation condition interaction remained significant, with greater micro-offline gains and lower micro-online gains in performance in the active as compared to the control condition. After exclusion of the extreme participant, the interaction and the micro-offline effect remained significant, whereas the micro-online effect did not reach significance.

### 8. Supplemental macro timescale results

**Table S12. Between-experiment comparison of macro-offline changes in performance speed.**

| Effect / Comparison | Statistic |
|---|---|
| **A. Macro gains between experiments — 2 (Experiment) × 2 (Stimulation condition) mixed ANOVA** | |
| Experiment | $F(1.00, 62.00) = 1.28$, $p = .263$, $\eta^2p = 0.020$ |
| Experiment × Stimulation condition | $F(1.00, 62.00) = 0.62$, $p = .432$, $\eta^2p = 0.010$ |

### 9. Protocol deviations

There were occasional issues that arose during data acquisition which resulted in deviations from the standard protocol. These deviations are reported in Table S13.

**Table S13. Protocol deviations.**

| Experiment | ID | Deviation |
|---|---|---|
| Rest-TIS | E6 | **(1)** During Visit 2 Training 2, the last 2 rest blocks (rest blocks #19 and #20) were not stimulated due to experimental error (experimenter interrupted the stimulation). The task was fully completed (20 blocks of practice in total, but only 18 rest blocks were stimulated). All task data were included in the analyses (including the 2 practice blocks #19 and #20 that preceded the non-stimulated rest epochs). |
| Rest-TIS | E8 | Visit 1: Actiwatch data corrupted. |
| Task-TIS | E9 | **(1)** During Visit 1 Training 1, the first 1.5 blocks were not stimulated due to experimental error (cable not plugged properly). The full session was restarted after fixing the issue which resulted in 1.5 unstimulated + 20 stimulated practice blocks in total. The data from the first 1.5 unstimulated blocks were not included in the analysis.<br>**(2)** During Visit 1 Training 2, due to computer malfunction, the task script crashed on the second to last rest block (i.e., rest block #19). The last block of practice (practice block #20) was therefore not practiced and not stimulated. The participant therefore only performed 19 blocks of practice during which stimulation was delivered (19 blocks practiced and stimulated). Data corresponding to the 19 blocks were included in the analyses. However, this participant was excluded from the Training phase x block repeated-measures ANOVA due to the missing block. Additionally, to ensure comparable summation of micro-online and micro-offline gains across visits, these gains were computed over the first 39 blocks (i.e., excluding the last block of Training 2) for both visits.<br>**(3)** During Visit 1 Test, due to computer malfunction, the task script crashed during the rest period preceding the last practice block #4. The script was restarted to complete the last block (4 blocks of practice in total, rest between blocks 3 and 4 was longer than by design due to task restart). All blocks were included in the analysis. |

| | | |
|---|---|---|
| | | **(4)** Actiwatch data missing for Visit 1 (participant forgot to wear the actiwatch). Sleep compliance was verified using the St. Mary's Hospital Sleep Questionnaire only. |
| Task-TIS | E13 | **(1)** During Visit 1 random SRTT, the script was restarted 4 times due to computer malfunction. The last 4 complete blocks of random SRTT were included in the analysis. |
| Rest-TIS | E20 | **(1)** During Visit 2 Training 2, the computer crashed in the middle of practice block #9. After script restart, the participant completed the remaining 11 blocks with rest stimulation. In total, the participant therefore performed 19.5 blocks of practice and received stimulation during 19 rest blocks (stimulation between practice blocks #9 and #10 was not delivered and the duration of this rest block was longer than by design due to script restart). All the data were included in the analyses (performance on practice block #9 includes less data than per design). |
| Task-TIS | E21 | **(1)** Visit 2: Due to a parameter entry error, the participant completed the spatial and motor tasks in the reversed order. **(2)** Actiwatch data missing for Visit 2 (participant forgot to wear the actiwatch). Sleep compliance was verified using the St. Mary's Hospital Sleep Questionnaire only. |
| Task-TIS | E25 | **(1)** Actiwatch data missing for Visit 2 (participant forgot to wear the actiwatch). Sleep compliance was verified using the St. Mary's Hospital Sleep Questionnaire only. |
| Task-TIS | E49 | **(1)** Actiwatch data missing for Visit 1 (participant forgot to wear the actiwatch). Sleep compliance was verified using the St. Mary's Hospital Sleep Questionnaire only. |
| Rest-TIS | E54 | **(1)** Visit 2: Blinding questionnaire data not logged by the experimenter. Participant was excluded from the corresponding statistical analyses. |
| Rest-TIS | E56 | **(1)** During Visit 1 Day 2 Retest, the participant accidentally pressed the Escape key at the beginning of Block 4, causing the task to terminate prematurely. The task was restarted approximately 1 minute later, and the participant completed an additional 1.5 blocks. Data from the 4 complete blocks were used in the analyses. |
| Task-TIS | E63 | **(1)** Visit 1: Actiwatch data corrupted/lost due to water damage. **(2)** Visit 2: Actiwatch data could not be retrieved due to device malfunction. |

### 10. Discarded trials (outliers and inaccurate)

**Table S14. Discarded trials for all task conditions, stimulation conditions and experiments.**

| Experiment | Condition | N | Total keys | Inaccurate | Outlier | Discarded |
|---|---|---|---|---|---|---|
| **A. Training (Blocks 1–40)** | | | | | | |
| task-TIS | Active | 32 | 118,077 | 4,061 (3.4%) | 2,359 (2.0%) | 6,420 (5.4%) |
| task-TIS | Control | 32 | 118,607 | 3,926 (3.3%) | 2,428 (2.0%) | 6,354 (5.4%) |
| rest-TIS | Active | 32 | 121,767 | 3,842 (3.2%) | 2,507 (2.1%) | 6,349 (5.2%) |
| rest-TIS | Control | 32 | 124,556 | 4,341 (3.5%) | 2,357 (1.9%) | 6,698 (5.4%) |
| 2 (Stimulation condition) × 2 (Experiment) mixed ANOVA on discarded trials (%) –; Experiment: $F(1, 62) = 0.318$, $p = .575$, $\eta^2G = 0.0032$; Stimulation condition × Experiment: $F(1, 62) = 1.329$, $p = .253$, $\eta^2G = 0.0081$. | | | | | | |
| **B. Post-training test (Blocks 41–44)** | | | | | | |
| task-TIS | Active TIS | 32 | 15,039 | 398 (2.6%) | 290 (1.9%) | 688 (4.6%) |
| task-TIS | Control | 32 | 15,071 | 423 (2.8%) | 286 (1.9%) | 709 (4.7%) |
| rest-TIS | Active TIS | 32 | 15,424 | 358 (2.3%) | 259 (1.7%) | 617 (4.0%) |
| rest-TIS | Control | 32 | 15,932 | 487 (3.1%) | 261 (1.6%) | 748 (4.7%) |

| | | | | | | |
|---|---|---|---|---|---|---|
| 2 (Stimulation condition) × 2 (Experiment) mixed ANOVA on discarded trials (%) –Experiment: $F(1, 62) = 0.384$, $p = .538$, $\eta^2G = 0.0043$; Stimulation condition × Experiment: $F(1, 62) = 1.575$, $p = .214$, $\eta^2G = 0.0078$. | | | | | | |
| **C. Day 2 retest (Blocks 45–48)** | | | | | | |
| task-TIS | Active TIS | 32 | 14,721 | 318 (2.2%) | 260 (1.8%) | 578 (3.9%) |
| task-TIS | Control | 32 | 15,379 | 405 (2.6%) | 261 (1.7%) | 666 (4.3%) |
| rest-TIS | Active TIS | 32 | 15,648 | 309 (2.0%) | 261 (1.7%) | 570 (3.6%) |
| rest-TIS | Control | 32 | 16,356 | 377 (2.3%) | 227 (1.4%) | 604 (3.7%) |
| 2 (Stimulation condition) × 2 (Experiment) mixed ANOVA on discarded trials (%) –Experiment: $F(1, 62) = 1.935$, $p = .169$, $\eta^2G = 0.0190$; Stimulation condition × Experiment: $F(1, 62) = 0.093$, $p = .762$, $\eta^2G = 0.0006$. | | | | | | |
| **D. Spatial transfer (4 blocks)** | | | | | | |
| task-TIS | Active TIS | 32 | 10,831 | 321 (3.0%) | 224 (2.1%) | 545 (5.0%) |
| task-TIS | Control | 32 | 11,126 | 478 (4.3%) | 232 (2.1%) | 710 (6.4%) |
| rest-TIS | Active TIS | 32 | 11,619 | 351 (3.0%) | 239 (2.1%) | 590 (5.1%) |
| rest-TIS | Control | 32 | 11,642 | 427 (3.7%) | 223 (1.9%) | 650 (5.6%) |
| 2 (Stimulation condition) × 2 (Experiment) mixed ANOVA on discarded trials (%) –Experiment: $F(1, 62) = 0.398$, $p = .531$, $\eta^2G = 0.0032$; Stimulation condition × Experiment: $F(1, 62) = 0.623$, $p = .433$, $\eta^2G = 0.0050$. | | | | | | |
| **E. Motor transfer (4 blocks)** | | | | | | |
| task-TIS | Active TIS | 32 | 11,399 | 378 (3.3%) | 226 (2.0%) | 604 (5.3%) |
| task-TIS | Control | 32 | 11,569 | 412 (3.6%) | 255 (2.2%) | 667 (5.8%) |
| rest-TIS | Active TIS | 32 | 12,885 | 366 (2.8%) | 222 (1.7%) | 588 (4.6%) |
| rest-TIS | Control | 32 | 12,772 | 468 (3.7%) | 224 (1.8%) | 692 (5.4%) |
| 2 (Stimulation condition) × 2 (Experiment) mixed ANOVA on discarded trials (%) –Experiment: $F(1, 62) = 2.465$, $p = .121$, $\eta^2G = 0.0197$; Stimulation condition × Experiment: $F(1, 62) = 0.505$, $p = .480$, $\eta^2G = 0.0040$. | | | | | | |
| **F. Novel-sequence transfer (4 blocks)** | | | | | | |
| task-TIS | Active TIS | 32 | 10,457 | 628 (6.0%) | 226 (2.2%) | 854 (8.2%) |
| task-TIS | Control | 32 | 10,295 | 480 (4.7%) | 206 (2.0%) | 686 (6.7%) |
| rest-TIS | Active TIS | 32 | 10,934 | 463 (4.2%) | 223 (2.0%) | 686 (6.3%) |
| rest-TIS | Control | 32 | 11,145 | 530 (4.8%) | 215 (1.9%) | 745 (6.7%) |
| 2 (Stimulation condition) × 2 (Experiment) mixed ANOVA on discarded trials (%) –Experiment: $F(1, 62) = 1.618$, $p = .208$, $\eta^2G = 0.0179$; Stimulation condition × Experiment: **$F(1, 62) = 7.366$, $p = .009$, $\eta^2G = 0.0345$.** Follow-up tests on the interaction – Active vs. Control: task-TIS $t(31) = 2.678$, $p = .012$, $dz = 0.47$; rest-TIS $t(31) = −1.083$, $p = .287$, $dz = −0.19$. task-TIS vs. rest-TIS: Active $t(62) = 2.571$, $p = .013$, $d = 0.64$; Control $t(62) = −0.422$, $p = .675$, $d = −0.11$. | | | | | | |

*Note.* Discarded trials include inaccurate transitions (incorrect key presses) and outlier inter-response intervals (individual IRIs exceeding the participant's block mean ± 3 SD). N = number of participants per cell. Percentages are relative to the total number of keypresses in that cell. Statistical tests: 2 (Experiment, between) × 2 (Stimulation condition, within) mixed ANOVAs on the percentage of discarded trials. Results of these ANOVAs (main effect of Stimulation condition, main effect of Experiment, and their interaction) are reported in the table below each panel (A–F). Significant interactions were followed up with paired t-tests (Active vs. Control within each experiment; Cohen's dz) and independent-samples t-tests (task-TIS vs. rest-TIS within each stimulation condition; Cohen's d).

### 11. Control analyses

#### Table S15. Baseline performance on the random SRTT: speed (A) and accuracy (B).

| Effect | F, df, p, $\eta^2p$ |
|---|---|

| | |
|---|---|
| **A. Baseline speed (mean RT on the random SRTT) — 4 (Block) × 2 (Stimulation condition)** | |
| **task-TIS (N = 32)** | |
| Block | **F(2.24, 69.31) = 8.84, p < .001, $\eta^2p$ = 0.222** |
| Stimulation condition | F(1.00, 31.00) = 0.23, p = .635, $\eta^2p$ = 0.007 |
| Block × Stimulation condition | F(2.17, 67.25) = 2.03, p = .135, $\eta^2p$ = 0.062 |
| **rest-TIS (N = 32)** | |
| Block | **F(2.60, 80.44) = 23.03, p < .001, $\eta^2p$ = 0.426** |
| Stimulation condition | F(1.00, 31.00) = 1.19, p = .285, $\eta^2p$ = 0.037 |
| Block × Stimulation condition | F(2.14, 66.40) = 1.79, p = .172, $\eta^2p$ = 0.055 |
| **Between experiments (N = 64)** | |
| Experiment | F(1.00, 62.00) = 3.03, p = .087, $\eta^2p$ = 0.047 |
| Experiment × Block | F(2.37, 146.81) = 1.31, p = .274, $\eta^2p$ = 0.021 |
| Experiment × Stimulation condition | F(1.00, 62.00) = 0.94, p = .337, $\eta^2p$ = 0.015 |
| Experiment × Block × Stimulation condition | **F(2.38, 147.71) = 3.07, p = .041, $\eta^2p$ = 0.047** |
| **B. Baseline accuracy (% correct responses on the random SRTT) — 4 (Block) × 2 (Stimulation condition)** | |
| **task-TIS (N = 32)** | |
| Block | F(2.84, 88.17) = 0.22, p = .870, $\eta^2p$ = 0.007 |
| Stimulation condition | F(1.00, 31.00) = 0.15, p = .700, $\eta^2p$ = 0.005 |
| Block × Stimulation condition | F(2.83, 87.71) = 0.29, p = .824, $\eta^2p$ = 0.009 |
| **rest-TIS (N = 32)** | |
| Block | F(2.15, 66.81) = 0.71, p = .503, $\eta^2p$ = 0.023 |
| Stimulation condition | F(1.00, 31.00) = 0.21, p = .646, $\eta^2p$ = 0.007 |
| Block × Stimulation condition | F(2.75, 85.19) = 1.16, p = .329, $\eta^2p$ = 0.036 |
| **Between experiments (N = 64)** | |
| Experiment | F(1.00, 62.00) = 2.13, p = .149, $\eta^2p$ = 0.033 |
| Experiment × Block | F(2.77, 171.66) = 0.38, p = .751, $\eta^2p$ = 0.006 |
| Experiment × Stimulation condition | F(1.00, 62.00) = 0.37, p = .547, $\eta^2p$ = 0.006 |
| Experiment × Block × Stimulation condition | F(2.85, 176.50) = 0.85, p = .465, $\eta^2p$ = 0.013 |

*Note.* Greenhouse–Geisser correction applied to within-subject factors with more than one numerator df. Mean response time and accuracy on the pseudo-random SRTT, performed before training under each stimulation condition, were analyzed with 4 (Block) × 2 (Stimulation condition) repeated-measures ANOVAs within each experiment and with 2 (Experiment) × 4 (Block) × 2 (Stimulation condition) mixed ANOVAs between experiments. Baseline speed and accuracy did not differ between stimulation conditions in either experiment. The Experiment × Block × Stimulation condition interaction on speed was not supported by follow-up analyses, as the Block × Stimulation condition interaction was not significant in either experiment. Significant effects (p < .05) are indicated in bold.

### Table S16. Performance speed (A-B) and accuracy (C-D) per sequence type (Set A vs. Set B).

| Effect | F, df, p, $\eta^2p$ |
|---|---|
| **A. Training speed — 2 (Training phase) × 20 (Block) × 2 (Set)** | |
| **task-TIS (N = 32)** | |
| Training phase | **F(1.00, 31.00) = 163.94, p < .001, $\eta^2p$ = 0.841** |
| Block | **F(3.59, 111.35) = 77.50, p < .001, $\eta^2p$ = 0.714** |
| Set | **F(1.00, 31.00) = 11.66, p = .002, $\eta^2p$ = 0.273** |
| Training phase × Block | **F(4.30, 133.36) = 71.74, p < .001, $\eta^2p$ = 0.698** |

| | |
|---|---|
| Training phase × Set | **F(1.00, 31.00) = 5.86, p = .022, $\eta^2p$ = 0.159** |
| Block × Set | **F(4.47, 138.65) = 2.87, p = .021, $\eta^2p$ = 0.085** |
| Training phase × Block × Set | F(5.46, 169.15) = 2.01, p = .073, $\eta^2p$ = 0.061 |
| **rest-TIS (N = 32)** | |
| Training phase | **F(1.00, 31.00) = 170.82, p < .001, $\eta^2p$ = 0.846** |
| Block | **F(3.49, 108.25) = 80.86, p < .001, $\eta^2p$ = 0.723** |
| Set | F(1.00, 31.00) = 1.51, p = .228, $\eta^2p$ = 0.046 |
| Training phase × Block | **F(4.54, 140.62) = 82.42, p < .001, $\eta^2p$ = 0.727** |
| Training phase × Set | F(1.00, 31.00) = 0.49, p = .490, $\eta^2p$ = 0.016 |
| Block × Set | F(3.99, 123.55) = 0.60, p = .663, $\eta^2p$ = 0.019 |
| Training phase × Block × Set | F(3.66, 113.45) = 0.91, p = .456, $\eta^2p$ = 0.029 |
| **Between experiments (N = 64)** | |
| Experiment | F(1.00, 62.00) = 0.67, p = .416, $\eta^2p$ = 0.011 |
| Experiment × Training phase | F(1.00, 62.00) = 0.13, p = .724, $\eta^2p$ = 0.002 |
| Experiment × Block | F(3.76, 233.27) = 0.65, p = .620, $\eta^2p$ = 0.010 |
| Experiment × Set | F(1.00, 62.00) = 2.93, p = .092, $\eta^2p$ = 0.045 |
| Experiment × Training phase × Block × Set | F(4.95, 307.19) = 1.76, p = .122, $\eta^2p$ = 0.028 |
| **B. Post-training test speed — 4 (Block) × 2 (Set)** | |
| **task-TIS (N = 32)** | |
| Block | F(2.50, 77.64) = 0.44, p = .693, $\eta^2p$ = 0.014 |
| Set | **F(1.00, 31.00) = 7.89, p = .009, $\eta^2p$ = 0.203** |
| Block × Set | F(1.86, 57.56) = 2.04, p = .142, $\eta^2p$ = 0.062 |
| **rest-TIS (N = 32)** | |
| Block | F(2.57, 79.70) = 1.66, p = .189, $\eta^2p$ = 0.051 |
| Set | F(1.00, 31.00) = 1.84, p = .185, $\eta^2p$ = 0.056 |
| Block × Set | F(2.41, 74.58) = 0.12, p = .915, $\eta^2p$ = 0.004 |
| **Between experiments (N = 64)** | |
| Experiment | F(1.00, 62.00) = 0.89, p = .350, $\eta^2p$ = 0.014 |
| Experiment × Block | F(2.55, 158.02) = 0.02, p = .991, $\eta^2p$ = 0.000 |
| Experiment × Set | F(1.00, 62.00) = 0.62, p = .434, $\eta^2p$ = 0.010 |
| Experiment × Block × Set | F(2.37, 146.82) = 1.00, p = .381, $\eta^2p$ = 0.016 |
| **C. Training accuracy — 2 (Training phase) × 20 (Block) × 2 (Set)** | |
| **task-TIS (N = 32)** | |
| Training phase | **F(1.00, 31.00) = 20.53, p < .001, $\eta^2p$ = 0.398** |
| Block | F(10.60, 328.65) = 0.96, p = .481, $\eta^2p$ = 0.030 |
| Set | **F(1.00, 31.00) = 21.50, p < .001, $\eta^2p$ = 0.410** |
| Training phase × Block | F(10.23, 317.20) = 1.71, p = .075, $\eta^2p$ = 0.052 |
| Training phase × Set | F(1.00, 31.00) = 0.56, p = .458, $\eta^2p$ = 0.018 |
| Block × Set | F(10.67, 330.64) = 1.25, p = .254, $\eta^2p$ = 0.039 |
| Training phase × Block × Set | F(10.72, 332.26) = 1.23, p = .270, $\eta^2p$ = 0.038 |
| **rest-TIS (N = 32)** | |
| Training phase | **F(1.00, 31.00) = 13.96, p < .001, $\eta^2p$ = 0.310** |
| Block | F(7.66, 237.59) = 1.65, p = .115, $\eta^2p$ = 0.051 |
| Set | **F(1.00, 31.00) = 5.18, p = .030, $\eta^2p$ = 0.143** |

| | |
|---|---|
| Training phase × Block | F(8.73, 270.59) = 1.13, p = .342, $\eta^2p$ = 0.035 |
| Training phase × Set | **F(1.00, 31.00) = 5.71, p = .023, $\eta^2p$ = 0.156** |
| Block × Set | F(8.73, 270.53) = 0.61, p = .781, $\eta^2p$ = 0.019 |
| Training phase × Block × Set | F(8.42, 261.11) = 1.29, p = .244, $\eta^2p$ = 0.040 |
| **Between experiments (N = 64)** | |
| Experiment | F(1.00, 62.00) = 0.09, p = .769, $\eta^2p$ = 0.001 |
| Experiment × Training phase | F(1.00, 62.00) = 0.88, p = .353, $\eta^2p$ = 0.014 |
| Experiment × Block | F(11.15, 691.56) = 0.70, p = .739, $\eta^2p$ = 0.011 |
| Experiment × Set | F(1.00, 62.00) = 0.15, p = .700, $\eta^2p$ = 0.002 |
| Experiment × Training phase × Block × Set | F(13.08, 810.74) = 1.43, p = .138, $\eta^2p$ = 0.023 |
| **D. Post-training test accuracy — 4 (Block) × 2 (Set)** | |
| **task-TIS (N = 32)** | |
| Block | F(2.30, 71.41) = 2.67, p = .069, $\eta^2p$ = 0.079 |
| Set | **F(1.00, 31.00) = 7.44, p = .010, $\eta^2p$ = 0.193** |
| Block × Set | **F(2.19, 68.02) = 5.49, p = .005, $\eta^2p$ = 0.150** |
| **rest-TIS (N = 32)** | |
| Block | F(2.76, 85.60) = 0.90, p = .440, $\eta^2p$ = 0.028 |
| Set | F(1.00, 31.00) = 0.15, p = .698, $\eta^2p$ = 0.005 |
| Block × Set | F(2.69, 83.52) = 1.57, p = .206, $\eta^2p$ = 0.048 |
| **Between experiments (N = 64)** | |
| Experiment | F(1.00, 62.00) = 0.00, p = .991, $\eta^2p$ = 0.000 |
| Experiment × Block | **F(2.57, 159.21) = 3.30, p = .028, $\eta^2p$ = 0.051** |
| Experiment × Set | F(1.00, 62.00) = 3.26, p = .076, $\eta^2p$ = 0.050 |
| Experiment × Block × Set | F(2.71, 167.98) = 2.37, p = .078, $\eta^2p$ = 0.037 |

*Note.* Greenhouse–Geisser correction applied to within-subject factors with more than one numerator df. Set = sequence type (Set A vs. Set B), collapsed across stimulation conditions. Speed (cIRI, in s) and accuracy (% correct transitions) during training were analyzed with 2 (Training phase) × 20 (Block) × 2 (Set) repeated-measures ANOVAs and during the post-training test with 4 (Block) × 2 (Set) ANOVAs, within each experiment and with the corresponding mixed ANOVAs including Experiment as a between-subjects factor. Effects of Set reflect differences in sequence difficulty (i.e., sequence A was overall performed slower and less accurately than sequence B; Sequence A: 41321342; Sequence B: 23142143) that are orthogonal to the stimulation manipulation, as sets were counterbalanced across conditions. Significant effects (p < .05) are indicated in bold.

#### Table S17. Order effects (Visit 1 vs. Visit 2) on performance speed (A–B) and accuracy (C–D).

| Effect | F, df, p, $\eta^2p$ |
|---|---|
| **A. Training speed — 2 (Training phase) × 20 (Block) × 2 (Visit)** | |
| **task-TIS (N = 32)** | |
| Training phase | **F(1.00, 31.00) = 163.94, p < .001, $\eta^2p$ = 0.841** |
| Block | **F(3.59, 111.35) = 77.50, p < .001, $\eta^2p$ = 0.714** |
| Visit | **F(1.00, 31.00) = 12.43, p = .001, $\eta^2p$ = 0.286** |
| Training phase × Block | **F(4.30, 133.36) = 71.74, p < .001, $\eta^2p$ = 0.698** |
| Training phase × Visit | **F(1.00, 31.00) = 24.27, p < .001, $\eta^2p$ = 0.439** |
| Block × Visit | **F(5.27, 163.31) = 8.26, p < .001, $\eta^2p$ = 0.210** |
| Training phase × Block × Visit | **F(5.84, 181.15) = 4.68, p < .001, $\eta^2p$ = 0.131** |
| **rest-TIS (N = 32)** | |

| | |
|---|---|
| Training phase | **$F(1.00, 31.00) = 170.82$, $p < .001$, $\eta^2p = 0.846$** |
| Block | **$F(3.49, 108.25) = 80.86$, $p < .001$, $\eta^2p = 0.723$** |
| Visit | **$F(1.00, 31.00) = 7.73$, $p = .009$, $\eta^2p = 0.200$** |
| Training phase × Block | **$F(4.54, 140.62) = 82.42$, $p < .001$, $\eta^2p = 0.727$** |
| Training phase × Visit | **$F(1.00, 31.00) = 24.34$, $p < .001$, $\eta^2p = 0.440$** |
| Block × Visit | **$F(5.40, 167.49) = 10.17$, $p < .001$, $\eta^2p = 0.247$** |
| Training phase × Block × Visit | **$F(4.78, 148.21) = 6.96$, $p < .001$, $\eta^2p = 0.183$** |
| **Between experiments (N = 64)** | |
| Experiment | $F(1.00, 62.00) = 0.67$, $p = .416$, $\eta^2p = 0.011$ |
| Experiment × Training phase | $F(1.00, 62.00) = 0.13$, $p = .724$, $\eta^2p = 0.002$ |
| Experiment × Block | $F(3.76, 233.27) = 0.65$, $p = .620$, $\eta^2p = 0.010$ |
| Experiment × Visit | $F(1.00, 62.00) = 0.86$, $p = .357$, $\eta^2p = 0.014$ |
| Experiment × Training phase × Block × Visit | $F(6.28, 389.54) = 0.52$, $p = .799$, $\eta^2p = 0.008$ |
| **B. Post-training test speed — 4 (Block) × 2 (Visit)** | |
| **task-TIS (N = 32)** | |
| Block | $F(2.50, 77.64) = 0.44$, $p = .693$, $\eta^2p = 0.014$ |
| Visit | $F(1.00, 31.00) = 1.72$, $p = .199$, $\eta^2p = 0.053$ |
| Block × Visit | $F(1.91, 59.21) = 0.93$, $p = .396$, $\eta^2p = 0.029$ |
| **rest-TIS (N = 32)** | |
| Block | $F(2.57, 79.70) = 1.66$, $p = .189$, $\eta^2p = 0.051$ |
| Visit | $F(1.00, 31.00) = 0.08$, $p = .775$, $\eta^2p = 0.003$ |
| Block × Visit | $F(2.45, 76.01) = 1.11$, $p = .345$, $\eta^2p = 0.034$ |
| **Between experiments (N = 64)** | |
| Experiment | $F(1.00, 62.00) = 0.89$, $p = .350$, $\eta^2p = 0.014$ |
| Experiment × Block | $F(2.55, 158.02) = 0.02$, $p = .991$, $\eta^2p = 0.000$ |
| Experiment × Visit | $F(1.00, 62.00) = 0.44$, $p = .509$, $\eta^2p = 0.007$ |
| Experiment × Block × Visit | $F(2.40, 148.82) = 1.40$, $p = .249$, $\eta^2p = 0.022$ |
| **C. Training accuracy — 2 (Training phase) × 20 (Block) × 2 (Visit)** | |
| **task-TIS (N = 32)** | |
| Training phase | **$F(1.00, 31.00) = 20.53$, $p < .001$, $\eta^2p = 0.398$** |
| Block | $F(10.60, 328.65) = 0.96$, $p = .481$, $\eta^2p = 0.030$ |
| Visit | $F(1.00, 31.00) = 0.15$, $p = .703$, $\eta^2p = 0.005$ |
| Training phase × Block | $F(10.23, 317.20) = 1.71$, $p = .075$, $\eta^2p = 0.052$ |
| Training phase × Visit | $F(1.00, 31.00) = 2.47$, $p = .126$, $\eta^2p = 0.074$ |
| Block × Visit | $F(10.46, 324.23) = 0.85$, $p = .585$, $\eta^2p = 0.027$ |
| Training phase × Block × Visit | $F(10.82, 335.48) = 0.78$, $p = .656$, $\eta^2p = 0.025$ |
| **rest-TIS (N = 32)** | |
| Training phase | **$F(1.00, 31.00) = 13.96$, $p < .001$, $\eta^2p = 0.310$** |
| Block | $F(7.66, 237.59) = 1.65$, $p = .115$, $\eta^2p = 0.051$ |
| Visit | $F(1.00, 31.00) = 1.56$, $p = .221$, $\eta^2p = 0.048$ |
| Training phase × Block | $F(8.73, 270.59) = 1.13$, $p = .342$, $\eta^2p = 0.035$ |
| Training phase × Visit | $F(1.00, 31.00) = 0.00$, $p = .958$, $\eta^2p = 0.000$ |
| Block × Visit | $F(9.10, 281.97) = 1.22$, $p = .279$, $\eta^2p = 0.038$ |
| Training phase × Block × Visit | $F(8.36, 259.11) = 0.92$, $p = .499$, $\eta^2p = 0.029$ |

| | |
|---|---|
| **Between experiments (N = 64)** | |
| Experiment | $F(1.00, 62.00) = 0.09, p = .769, \eta^2p = 0.001$ |
| Experiment × Training phase | $F(1.00, 62.00) = 0.88, p = .353, \eta^2p = 0.014$ |
| Experiment × Block | $F(11.15, 691.56) = 0.70, p = .739, \eta^2p = 0.011$ |
| Experiment × Visit | $F(1.00, 62.00) = 0.62, p = .434, \eta^2p = 0.010$ |
| Experiment × Training phase × Block × Visit | $F(13.03, 807.71) = 0.83, p = .628, \eta^2p = 0.013$ |
| **D. Post-training test accuracy — 4 (Block) × 2 (Visit)** | |
| **task-TIS (N = 32)** | |
| Block | $F(2.30, 71.41) = 2.67, p = .069, \eta^2p = 0.079$ |
| Visit | $F(1.00, 31.00) = 0.23, p = .635, \eta^2p = 0.007$ |
| Block × Visit | $F(1.98, 61.51) = 0.05, p = .951, \eta^2p = 0.002$ |
| **rest-TIS (N = 32)** | |
| Block | $F(2.76, 85.60) = 0.90, p = .440, \eta^2p = 0.028$ |
| Visit | **$F(1.00, 31.00) = 4.56, p = .041, \eta^2p = 0.128$** |
| Block × Visit | $F(2.66, 82.45) = 0.24, p = .846, \eta^2p = 0.008$ |
| **Between experiments (N = 64)** | |
| Experiment | $F(1.00, 62.00) = 0.00, p = .991, \eta^2p = 0.000$ |
| Experiment × Block | **$F(2.57, 159.21) = 3.30, p = .028, \eta^2p = 0.051$** |
| Experiment × Visit | $F(1.00, 62.00) = 3.96, p = .051, \eta^2p = 0.060$ |
| Experiment × Block × Visit | $F(2.60, 161.25) = 0.11, p = .939, \eta^2p = 0.002$ |

*Note.* Greenhouse–Geisser correction applied to within-subject factors with more than one numerator df. Visit = experimental visit (Visit 1 vs. Visit 2), collapsed across stimulation conditions. Speed (cIRI, in s) and accuracy (% correct transitions) during training were analyzed with 2 (Training phase) × 20 (Block) × 2 (Visit) repeated-measures ANOVAs and during the post-training test with 4 (Block) × 2 (Visit) ANOVAs, within each experiment and with the corresponding mixed ANOVAs including Experiment as a between-subjects factor. Effects involving the visit factor reflect general practice effects across visits that are orthogonal to the stimulation manipulation, as condition order was counterbalanced. During training, performance speed was faster overall during the second visit; this practice advantage was most pronounced during the early blocks of Training 1 and progressively decreased with practice, as indicated by the interactions between Visit and Training phase/Block (A). These visit effects were no longer significant during the post-training test in either experiment, indicating that performance speed had reached comparable levels across visits by the end of training (B). For accuracy, no visit-related effects were observed during training (C); during the post-training test, accuracy was slightly higher during the second visit in the rest-TIS experiment only, with no visit effect in the task-TIS experiment or in the between-experiments analysis (D). Visit effects did not interact with Experiment in any analysis. Significant effects ($p < .05$) are indicated in bold.
